# Assessing Drought Resilience and Identification of High Yielding Upland Rice Varieties through Phenology, Growth and Yield Traits

**DOI:** 10.64898/2025.12.20.695743

**Authors:** Tajamul Hussain, Jakarat Anothai, Charassri Nualsri, Awais Ali, Thanet Khomphet

## Abstract

Drought stress is the major yield limiting factor in upland rice production where the moisture availability is highly variable. Understanding and evaluating how upland rice responds to drought stress is critical to improving resilience and yield stability. In this study performance of sixteen upland rice varieties were evaluated under non-stressed, moderately stressed and highly stressed conditions. Drought stress was introduced by irrigating upland rice at 70% and 50% field capacity (FC) whereas non-stress treatment was irrigated at 100% FC. Irrigation in moderately stressed and highly stressed conditions was also withheld for six days at lateral crop stages to observe temporary wilting by inducing a stress interval. Data on agronomic traits of upland rice was collected in three experimental replications. Results indicated that performance of upland rice varieties was significantly altered under stress conditions and highest performance was observed under non-stressed conditions. Yield losses for short duration and long duration varieties ranged 35-60% and 24-62% under moderate stress whereas it ranged 43-78% and 52-73% under highly stressed conditions, respectively. Overall varieties Dawk Kha, Khao^/^ Sai and Dawk Pa–yawm, indicated higher stability under stressed conditions therefore, these long duration varieties could be used for obtaining better yields under diverse agroclimatic conditions and under unpredicted weather patterns. Short duration Ma-led-nai-fai and long duration Goo Meung Lung and Bow Leb Nahag could be used for acquiring traits for higher tillering and panicle bearing capacity. Short heighted varieties such as Jao Daeng, Sahm Deuan and Ma-led-nai-fai could be used in breeding for short heighted new varieties to overcome lodging concerns. Strong significant association of GMP, STI, M_PRO_, M_HAR_ with grain yield under non-stressed, moderately stressed and highly stressed conditions indicated that these indices were appropriate for their use as selection criteria for drought resilience.

**Graphical Abstract:** 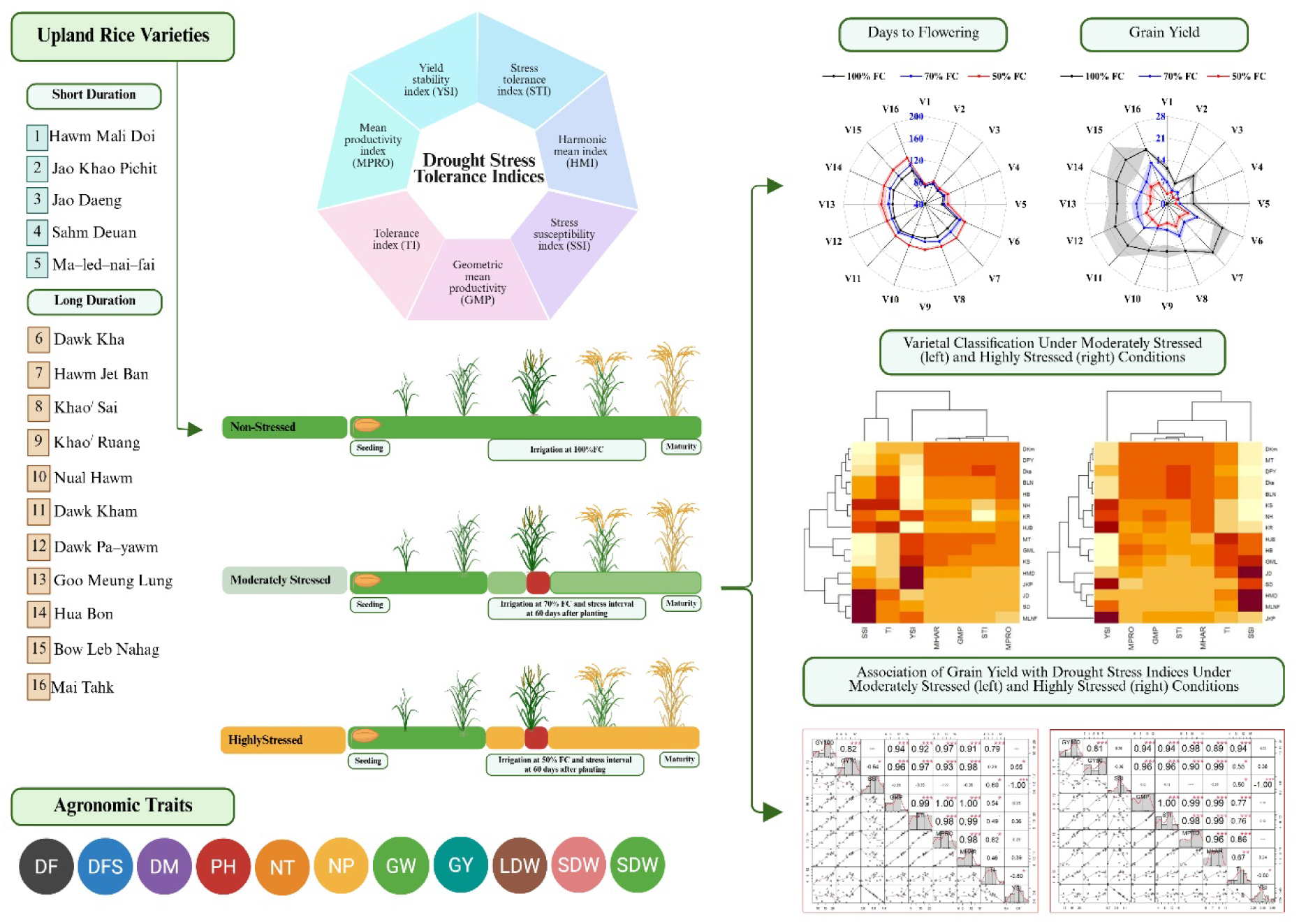

## 1. Introduction

Rice is one of the most vital food crops grown in more than 100 countries and serves as staple food for more than three billion people worldwide. Rice plays a central role in global food and nutritional security. Rice is cultivated in range of agroecosystems under variable agroclimatic conditions and management practices. Among abiotic stresses, drought remains the most critical environmental factor to rice production particularly in upland ecosystems (Yang et al., 2022) where the seasonal variability in rainfalls induce stress at critical crop growth stages and impacts rice productivity. Rice is highly sensitive to drought stress especially during its reproductive stages (Bhandari et al., 2023; Jiang et al., 2025; Liberatore et al., 2025). This sensitivity poses a serious threat to food security, a challenge exacerbated by climate change (Mansour et al., 2021; Yang et al., 2022). Increasing variability in seasonal rainfall patterns, extreme weather events and prolonged drought stress periods are severely threatening rice productivity (Rehmani et al., 2021; Fu et al., 2023; Joseph et al., 2023). Studies involving climate models consistently predicted that escalating drought severity will pose a critical challenge to global rice production (Chen et al., 2025; Liu et al., 2025). In this scenario, sustaining rice productivity in upland ecosystems is essential for meeting food security.

Drought stress affects upland rice performance across all growth stages however, the sensitivity to drought stress varies. The impact of drought stress on rice exhibits distinct physiological and yield responses (Hussain et al., 2018; Bhandari et al., 2023; Jarin et al., 2024). Drought occurrence at different crop stages significantly impairs leaf numbers and size, tillering capacity and stem elongation which affects panicle emergence and ultimately reduces grain yield (Zulkarnain et al., 2013; Alou et al., 2018; Khotasena et al., 2022). Drought at tillering stage can affect effective tillers that may influence overall canopy structure whereas drought stress at heading and flowering stage may significantly reduce grain numbers in panicles and seed setting (Jiang et al., 2025). Drought stress influence water regulation (Jiang et al., 2025) which impacts grain filling rate, grain numbers and grain weight (Bhandari et al., 2023) that ultimately influence final yields. Rice flowering is highly dependent on water availability. In contrast to other plant species such as *Arabidopsis*, drought in rice delays flowering (Rahman et al., 2002; Zhang et al., 2016) and maturity (Wopereis et al., 1996; Hussain et al., 2018). Stress occurrence causes pollen sterility, ovule abortion, reduces grain filling and grain weight leading to low rice productivity (Jarin et al., 2024; Jiang et al., 2025). Short and long duration varieties exhibit varying responses to drought stress. Short duration varieties may experience severe yield losses if drought occurs at lateral crop growth stages under natural conditions however, they are also used as drought avoiding varieties to escape drought stress. Long duration varieties have potential to recover from stress; however, yield losses are common. According to Ichsan et al. (Ichsan et al., 2020), stress tolerant varieties are capable of sustaining rice production. Drought tolerant varieties also exhibit smaller reduction in grain yield under drought stress compared to susceptible varieties (Hallajian et al., 2024; Sakran et al., 2025). Varieties maintaining final yields are considered tolerant and are preferred by growers. Variable varietal response to drought stress indicates the distinct impacts on plant’s physiological processes. Therefore, understanding and evaluating specific impacts of drought stress on varietal performance is critical for developing better management and crop improvement strategies for improved yields and stability (Habib et al., 2024) across different agroclimatic conditions.

To navigate the complexity in evaluating varietal performance under drought stress, researchers employ various techniques and drought assessment methods (Todaka et al., 2015; Singh et al., 2017; Hussain et al., 2021a; Hallajian et al., 2024; Sakran et al., 2025). Drought scoring methods are used to directly assess and select tolerant genotypes (IRRI, 2014; Ahmad et al., 2020; El-Refaee et al., 2023). Stress tolerance indices have been widely used to determine relationship between yield under non stressed and stressed conditions with different intensities and drought stress evaluation (Fischer and Maurer, 1978; Rosielle and Hamblin, 1981; Bouslama and Schapaugh, 1984; Clarke et al., 1984; Hossain et al., 1990; Fernandez, 1992; Schneider et al., 1997; Rashid et al., 2003; Anwar et al., 2011; Wasae, 2021; Mansour et al., 2021; Debbarma et al., 2024; Hallajian et al., 2024). Stress indices are dependent on stress tolerance or susceptibility of a variety (Clarke et al., 1984; Fernandez, 1992) and help elucidate the relationship between yield potential and yield stability under drought stress. Along with drought stress indices, relative yield is a measure to evaluate the stability or reduction in yields of a variety compared to tested population that determines stress tolerance or susceptibility of a variety. Drought stress indices include stress susceptibility index (SSI), geometric mean productivity (GMP), harmonic mean index (M_HAR_), mean productivity index (M_PRO_), yield stability index (YSI) and tolerance index (TI) (Fischer and Maurer, 1978; Rosielle and Hamblin, 1981; Bouslama and Schapaugh, 1984; Hossain et al., 1990; Fernandez, 1992; Schneider et al., 1997). Each of these indices provides a unique perspective on the trade-off between yield potential and stability of variety. Although these indices are widely used in drought stress evaluation, they share a key limitation that they are derived only from yield data and studies have demonstrated that strong positive and negative associations varied among these indices and grain yields (Golabadi et al., 2006; Arif et al., 2021; Hussain et al., 2021b). In this scenario, specific evaluations considering crop types, varieties and applied drought stress conditions are critical.

Considering the distinct varietal response of upland rice to drought stress and variable linkages among stress tolerance indices and grain yields, the objectives of current study, therefore, were to (i) evaluate agronomic performance of upland rice varieties under non-stressed, moderately stressed and highly stressed conditions and identify high yielding drought resilient varieties for recommendations and (ii) identify suitable drought stress indices for their application as reliable criteria in rice breeding to enhance drought resilience.

## 2. Material and Methods

### 2.1. Upland Rice Varieties

Five short duration upland rice varieties Hawm Mali Doi (HMD) [V1], Jao Khao Pichit (JKP) [V2], Jao Daeng (JD) [V31], Sahm Deuan (SD) [V4], Ma–led–nai–fai (MLNF) [V5] and eleven long duration upland rice varities including Dawk Kha (DKa) [V6], Hawm Jet Ban (HJB) [V7], Khao^/^ Sai (KS) [V8], Khao^/^ Ruang (KR) [V9], Nual Hawm (NH) [V10], Dawk Kham (DKm) [V11], Dawk Pa–yawm (DPY) [V12], Goo Meung Lung (GML) [V13], Hua Bon (HB) [V14], Bow Leb Nahag (BLN) [V15], and Mai Tahk (MT) [V16] were examined for drought stress resilience and susceptibility. Germplasm of these varieties was collected from various locations in Thailand. Characteristics of these varieties and the germplasm collection location are available in our previously published study (Hussain et al., 2023).

### 2.2. Experimental Setup and Management

The research experiment was conducted at Faculty of Natural Resources, Prince of Songkla University, Hat Yai, Thailand (7°00’14.6" N, 100°30’01.9" E). Coarse textured sandy clay loam field soil having 66.03% sand, 11.27 silt, and 22.70% of clay, was used for rice experiment. Soil contained 0.07 and 10.36% nitrogen and phosphorous that required optimal fertilization. Therefore, fertilizers were applied following an optimal rate of 75 kg N ha^-1^, 45 kg P ha^-1^ and 45 kg K^+^ ha^-1^ were applied. Upland rice varieties and drought stress intensities based on field capacity (FC) levels were experimental treatments in this study and varieties were tested under non-stressed (100% FC), moderately stressed (70% FC) and highly stressed (50% FC) conditions. To organize rice varieties that were unique in their maturity duration and facilitate irrigation management for experimental treatments the trial was established using strip plot design with three repetitions. Soil water contents were recorded to regulate stress treatments and irrigation applications. Planting was performed at 5 cm depth and twenty plants m^-2^ were maintained with 25 cm row and 20 cm planting distance. Plants were first irrigated uniformly till 60 days of planting to keep all plants under non-stress conditions and then treatments including non-stressed, moderately stressed and highly stressed at 100, 70 and 50% FC were employed. To induce stress interval irrigation was stopped at 68^th^ day of planting in short duration varieties and at 76^th^ day of planting in long duration varieties for moderately stressed and highly stressed treatments whereas irrigation in non-stress treatment continued without any alteration till harvest. Temporary wilting was observed in six days of stress induction following which the irrigation was resumed and moderately stressed and highly stressed treatments were maintained at 70% and 50% FC till maturity. Weeds were controlled manually.

### 2.3. Data Collection

Agronomic data for all upland rice varieties on days to first seed, flowering duration, maturity, plant height, number of tillers and number of panicles, grain weight, leaf, stem and total dry weights and grain yield were collected in all three repetitions. Plant height was measured from the stem base to the flag leaf tip or panicle. Number of days to first seed, flowering and maturity were recorded from planting to the date when 50% of total plant reached first seed, flowering and maturity, respectively. Number of tillers and panicles were counted to determine tillers and panicles for each variety. Selected plants were sampled, dried at 70 °C to a constant weight and dry weights were recorded.

### 2.4. Stress Tolerance Indices

Drought stress resilient and susceptible varieties were differentiated using drought stress tolerance indices. Yield stability index (Bouslama and Schapaugh, 1984) [1], stress tolerance index (Fernandez, 1992) [2], mean productivity index (Hossain et al., 1990) [3], tolerance index (Rosielle and Hamblin, 1981) [4], geometric mean productivity (Fernandez, 1992) [5], stress susceptibility index (Fischer and Maurer, 1978) [6] and harmonic mean index (Schneider et al., 1997) [7] were among the applied and tested drought stress indices in this assessment. Grain yield of all genotypes under non-stressed, moderately stressed and highly stressed conditions was used to determine relative yield performance under applied stressed intensities. Relative yield is a measure to determine the performance of a variety under applied drought stress, relative to other varieties being tested and it was computed by dividing the grain yield of each variety with the maximum observed yield under specific applied drought stress. Drought stress resilient verities were identified when they exhibited superior yield performance under non-stressed conditions, higher relative yield under stressed conditions and resulted promising values for drought stress indices. Highly correlated drought stressed indices were identified as promising indices for drought stress evaluation for upland rice varieties.

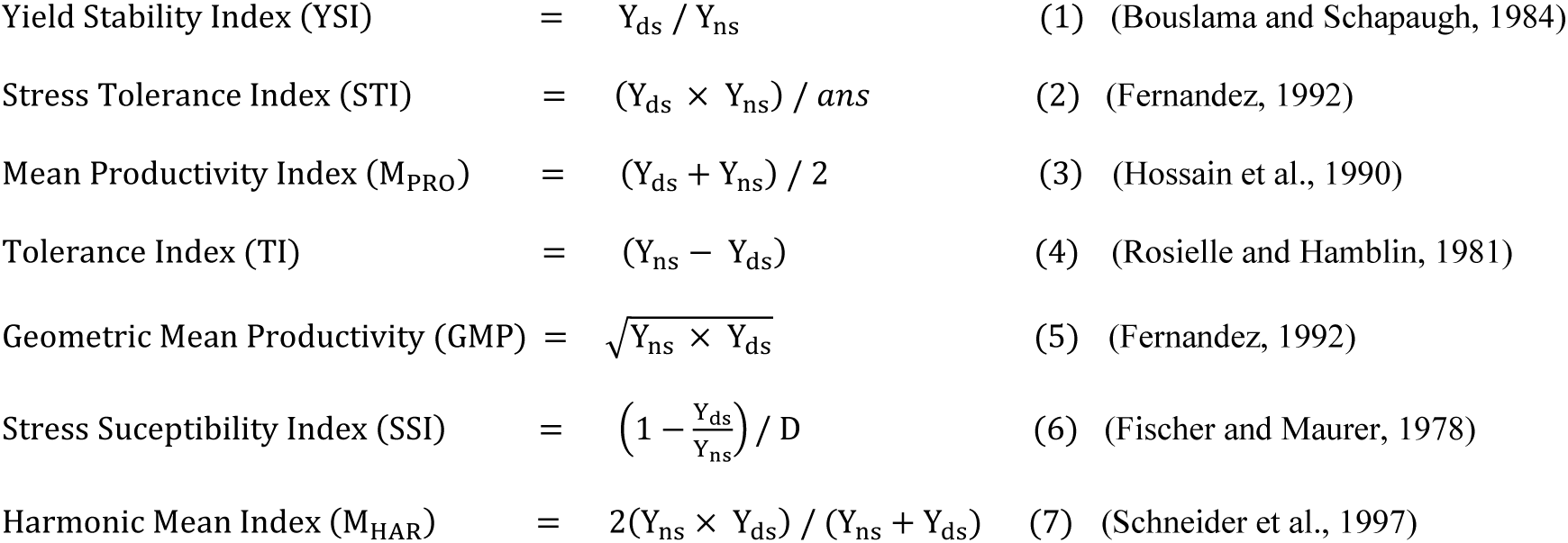

Where, Yds = mean grain yield under drought-stressed conditions, Yns = mean grain yield under non-stressed conditions, *D* = environmental stress intensity which is 1– (mean grain yield of variety under drought-stressed conditions / mean grain yield of genotypes under non-stressed conditions) and *ans* is average value for all examined varieties for grain yield under non-stressed conditions.

### 2.5. Statistical Analysis

Experimental data on agronomic traits was analyzed using Statistix 8.1 package (Analytical software, FL, USA) and analysis of variance (ANOVA) was conducted to determine the variety and treatment effects. Fisher’s least significant difference (LSD) method was used to compare means of studied traits and significance was considered at α ≤ 0.05. Radar (spider) plots were generated using Origin 2024b (Northampton, MA, USA) to illustrate the relative performance of agronomic traits of upland rice varieties under non-stressed, moderately stressed and highly stressed conditions. Pearson’s correlation coefficients were computed to determine association among agronomic traits of upland rice varieties and drought stress indices. R program (R Core Team, 2024) was used for visualization of correlation plots and matrix using “Corr” and “GGally” packages.

## 3. Results and Discussion

### 3.1. Upland rice performance to drought stress

Statistical analysis indicated that upland rice varieties significantly differed (*p* < 0.001) and were unique in their agronomic characteristics including days to flowering, days to first seed, days to maturity, plant height, number of tillers, number of panicles, 1000 grain weight, grain yield, leaf dry weight, stem dry weight and total dry matter (Table 1). Significant variation for varietal effect indicated the existing diversity in population. Diversity among upland rice varieties provide the options for breeding for example could be used for breeding specific traits in rice crop breeding program (Sun et al., 2022). Applied treatments also indicated highly significant (*p* < 0.001) response for all studied traits indicating that applied drought stress treatments significantly altered the performance of agronomic traits. Previous research has shown that drought stress significantly affected performance of agronomic traits of upland rice (Alou et al., 2018; Hussain et al., 2018; Sun et al., 2022; Hallajian et al., 2024; Yuwono et al., 2025). Findings from these indicated the sensitivity of rice to drought stress and suggested that drought occurrence can significantly impact rice production potential thus the screening for high yielding and drought resilient varieties is essential.

**Table 1:**
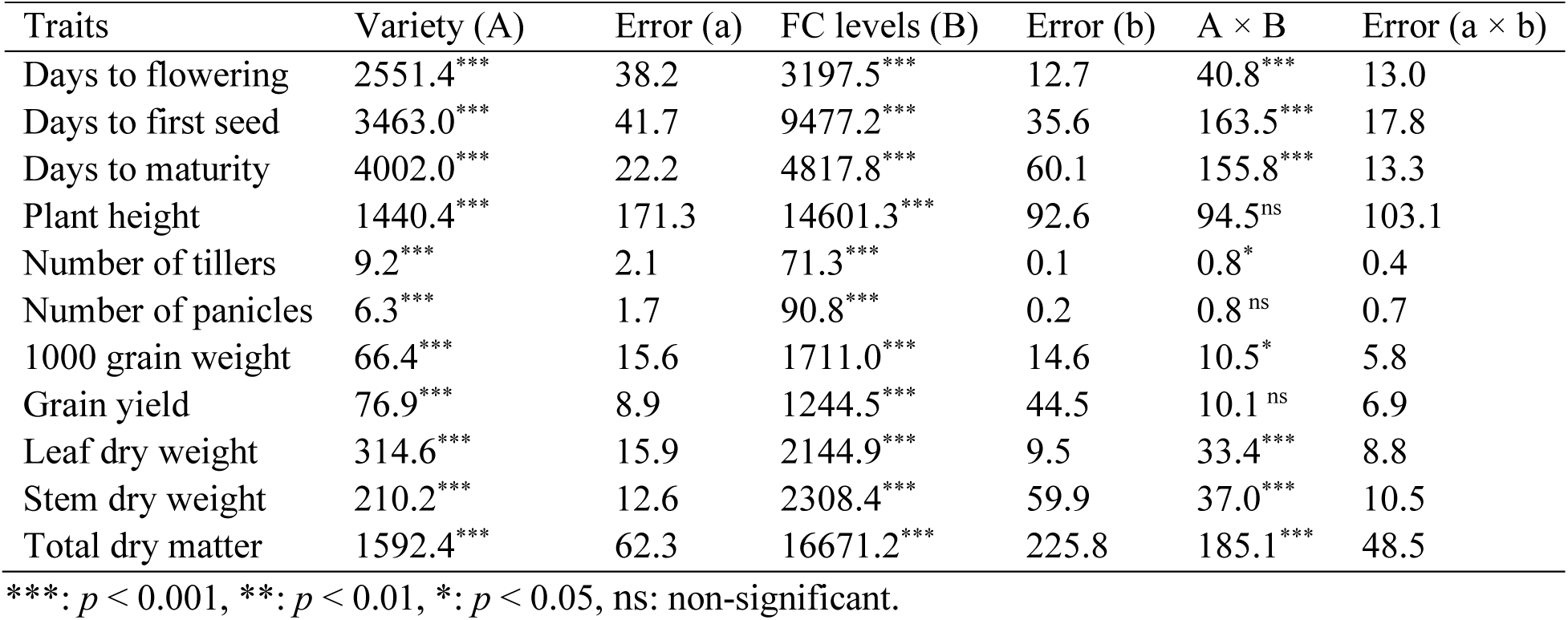
Mean squares of analysis of variance for agronomic traits of sixteen upland rice varieties observed under non-stressed, moderately stressed and highly stressed conditions (FC levels).

### 3.2. Phenological traits

Phenological traits including days to flowering, days to first seed and days to maturity were significantly affected under applied drought stress and varied among upland rice varieties (Figure 1a). Under non-stressed conditions, days to flowering ranged 72-80 and 102-108 days for short duration and long duration varieties, respectively. In this study drought stress delayed flowering in all upland rice varieties and days to flowering ranged 73-82 and 105-116 days for short duration and 76-88 and 120-131 for long duration varieties under moderately stressed and highly stressed conditions, respectively. The difference in increase in number days to flowering was higher in long duration varieties compared to short duration varieties under stressed conditions as it ranged 1-4 days and 3-10 days under moderately stressed conditions and ranged 4-12 days and 14-25 days under highly stressed conditions (Table S1). Delay in flowering can occur in rice under drought conditions. Kang and Futakuchi (Kang and Futakuchi, 2019), while working with 10 upland-adapted rice genotypes, *i.e.* eight interspecific *Oryza sativa* L. ∼ *Oryza glaberrima* Steud. lines and two *Oryza sativa* varieties—WAB56-104, the *Oryza sativa* parent of eight interspecific lines and IRAT 109, reported that flowering was delayed under drought stress conditions compared to wet which was likely due to impaired photosynthesis and resource allocation. Drought-induced stomatal closure reduces photosynthetic carbon assimilation limiting the supply of assimilates to the developing panicle, thereby halting or delaying flower initiation and growth. Delayed in flowering under drought stress in rice has been reported in previous studies (Zhang et al., 2016; Hussain, 2017; Alou et al., 2018; Adeboye et al., 2021; Jiang et al., 2025).

**Figure 1:**
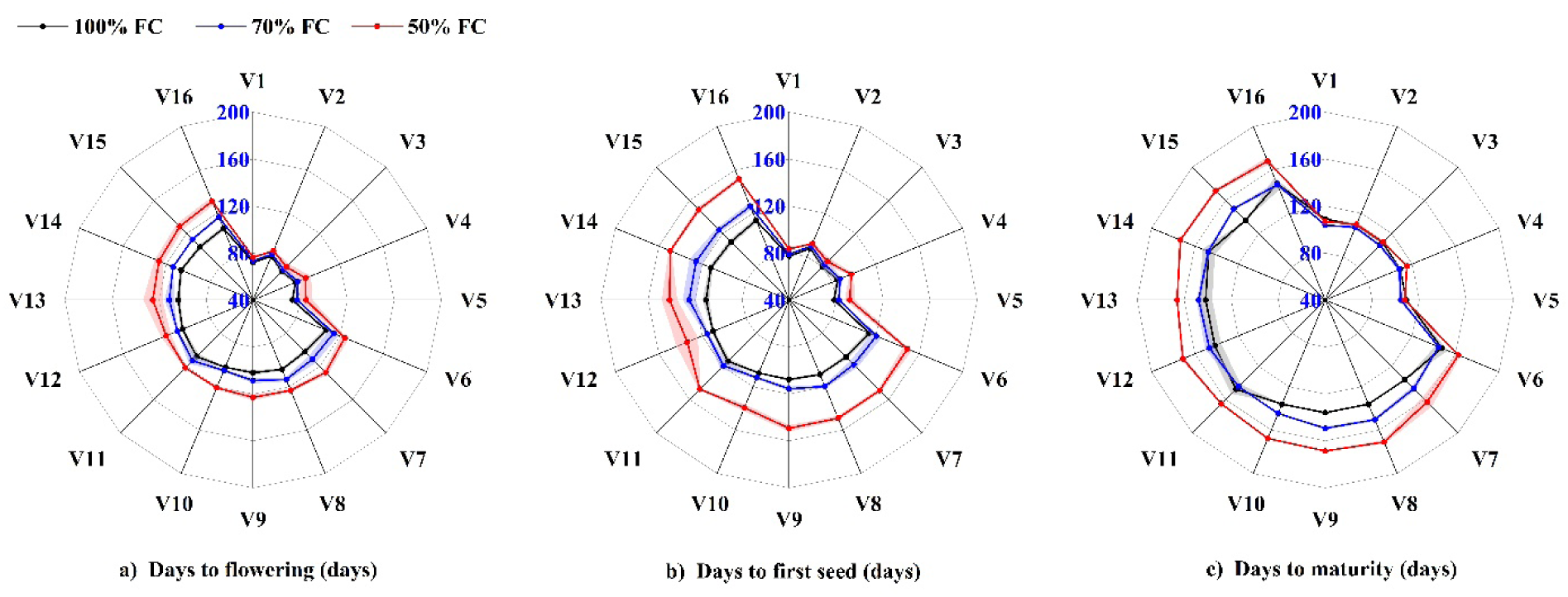
Days for flowering (DF), days to first seed (DS) and days to maturity (DM) of sixteen upland varieties under non-stressed (100%), moderately stressed (70%) and highly stressed (50%) conditions (field capacity levels). Shaded areas show ± standard errors for means (*n*=3).

Days to first seed (Figure 1b) under non-stressed conditions ranged 77-88 and 107-114 days for short duration and long duration varieties, respectively. Drought stress delayed formation of seed in all upland rice varieties and it ranged 79-90 and 112-126 days for short duration and 83-97 and 134-151 for long duration varieties under moderately stressed and highly stressed conditions, respectively. The difference in increase in number days to first seed was higher in long duration varieties compared to short duration varieties under stressed conditions as it ranged 2-4 days and 4-14 days under moderately stressed conditions and ranged 5-13 days and 24-42 days under highly stressed conditions (Table S1). These results align with the reported research that drought stress can significantly influence rice causing flowering disorders and decline in seed setting rate (Jiang et al., 2025).

Maturity duration (Figure 1c) indicated a variable trend compared to days to flowering and days to first seed. Short duration varieties matured earlier under moderately stressed and highly stressed conditions except for V1 and V5. Most long duration varieties matured earlier under moderately stressed and highly stressed conditions and the difference in delay ranged 5-14 days under moderately stressed and 15-36 days under highly stressed conditions. V14 matured at same time under non-stressed and moderately stressed conditions. Whereas V6, V11 and V16 matured earlier under moderately stressed conditions (Table S1). Late or early maturity of upland rice varieties under various applied stress conditions has been previously reported (Hussain, 2017; Pope et al., 2023). This is because under drought stress rice fails to grow well due to the physiological alterations, thereby delaying maturity. Some varieties mature early under stress conditions while others mature late. This is possibly due to different genetically programmed survival strategies to cope with water scarcity. These strategies fall primarily into two categories: drought escape (early maturity) and drought tolerance (late maturity). This variation in maturity duration provides the opportunity to use rice varieties for obtaining early or late maturity characteristics. Early maturity varieties could be used to benefit from drought escape. Chatterjee et al. (Chatterjee et al., 2024) who worked on the molecular insights in drought of upland rice, stated drought avoidance is a suitable strategy to combat drought stress as the varieties mature early avoiding stress conditions and maintaining their yield potential.

### 3.3. Growth and yield attributes

Plant height varied significantly among the short duration and long duration upland rice varieties (Figure 2a). Average plant height of short duration and long duration varieties ranged 124-162 cm and 158-183 cm under non stressed conditions. Drought stress reduced the overall plant height and for short and long duration varieties it ranged 99-145 and 139-164 under moderately stressed and ranged 97-136 and 124-150 cm under highly stressed conditions, respectively. Reduction in plant height ranged 9-20% and 13-22% for short duration and 8-21% and 15-30% for long duration varieties under moderately stressed and highly stressed conditions, respectively (Table S1). The reduction in plant height under drought stress is a mechanism that helps the plant conserve water and survive until conditions improve. Drought stress reduces cell turgidity which leads to reduced cell elongation resulting in stunted growth and overall shorter plant height; this morphological change is driven by a combination of physiological and hormonal responses (Sadhukhan et al., 2024). Reduction in plant height of upland rice varieties under drought stress is well documented in previous studies (Lanna et al., 2021; Jarin et al., 2024; Sagar et al., 2025).

**Figure 2:**
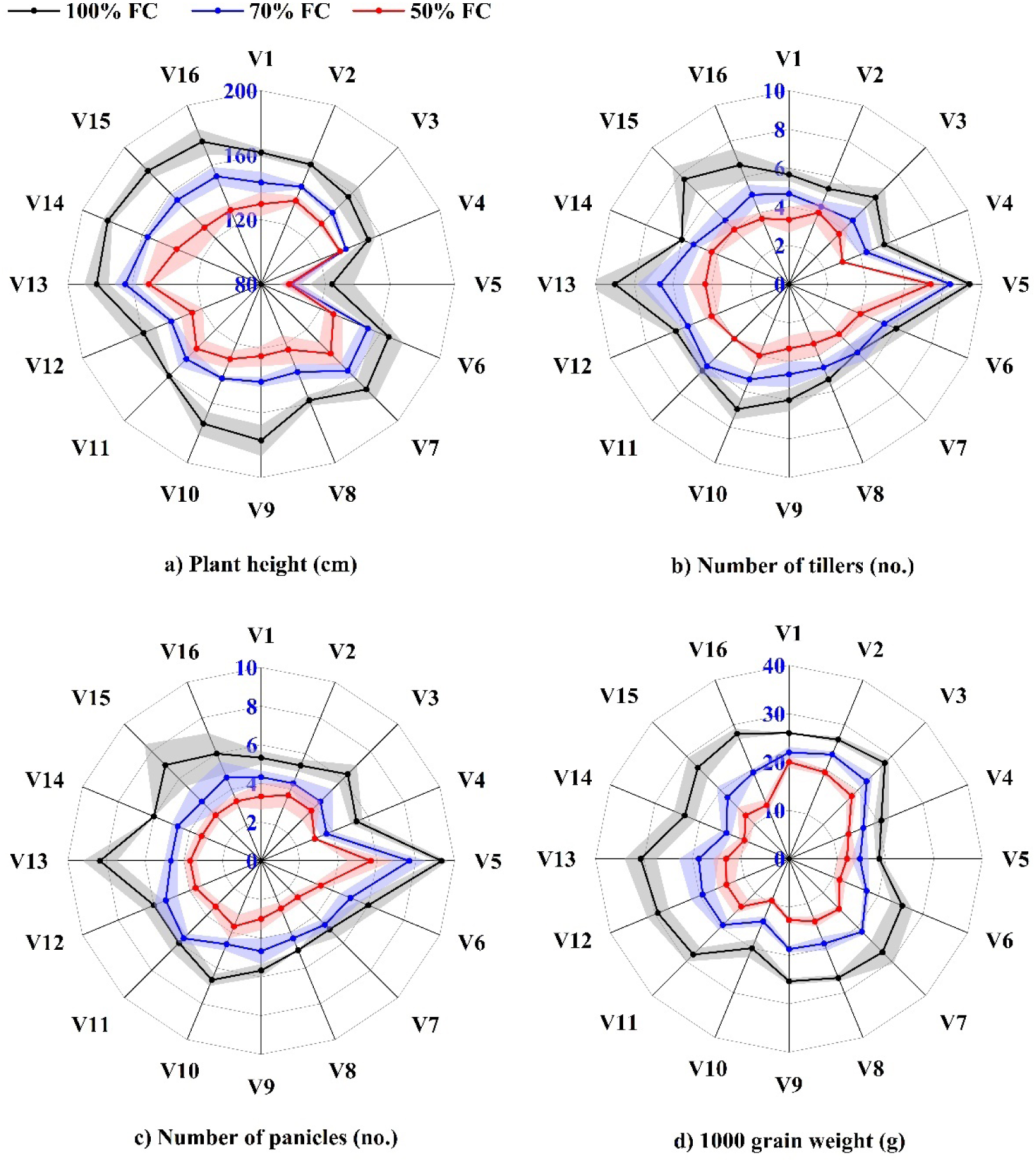
Plant height (PH), Number of tillers (NT), number of panicles (NP) and 1000 grain weight (1000GW) of sixteen upland varieties under non-stressed (100%), moderately stressed (70%) and highly stressed (50%) conditions (field capacity levels). Shaded areas show ± standard errors for means (*n*=3).

Number of tillers (Figure 2b, Table S1) were also affected under drought stress as some tillers failed to produce panicles. Upland rice varieties also differed in tillering capacity. Under non-stressed conditions number of tillers ranged 5-9 tillers per plant for both short and long duration varieties. However, under moderately stressed and highly stressed conditions average number of tillers per plant ranged 4-8 and 3-7 for short duration and ranged 5-7 and 3-4 for long duration varieties, respectively. Average number of tillers for V7 and V11 were not affected under moderately stressed conditions. Similar to number of tillers, upland rice varieties also differed in panicles bearing capacity (Figure 2c, Table S1). Under non-stressed conditions number of panicles ranged 5-9 panicles per plant for both short and long duration varieties. However, under moderately stressed and highly stressed conditions average number of panicles ranged 4-8 and 3-6 for short duration and ranged 4-6 and 3-4 for long duration varieties, respectively. Negative impact of drought stress on upland rice tillering and panicle bearing capacity has been reported in previous studies (Alou et al., 2018; Konaté et al., 2022). Under drought stress, rice plants fail to develop panicles leaving more tillers unproductive. This strong correlation between tillers and panicles confirms that higher tiller production supports a greater number of panicles. The outcome in the present study is consistent with previous research demonstrating that drought stress increases tiller mortality (Zain et al., 2014) and terminal-stage stress negatively impact panicle formation by altering source-sink dynamics (Davatgar et al., 2012).

Grain weight is useful indicator for determining overall yields. 1000 grain weight (Figure 2d) for short duration and long duration varieties ranged 19-28 and 20-31 grams (g), respectively under non-stressed conditions. Grain weight significantly reduced under drought conditions and ranged 15-23 and 12-20 g for short duration and ranged 14-21 and 9-15 g for long duration varieties under moderately stressed and highly stressed conditions, respectively. The difference in decrease in average grain weight was higher in long duration varieties compared to short duration varieties under stressed conditions as it ranged 13-21% and 22-40% under moderately stressed conditions and ranged 23-36% and 46-58% under highly stressed conditions (Table S2). Drought stress negatively impacts grain weight of rice varieties. The formation of underdeveloped, lighter grains, and thus reduced grain weight, is a primary consequence of drought stress disrupting essential physiological functions such as photosynthesis and nutrient uptake especially during the reproductive phase (da Mata et al., 2023; Sadhukhan et al., 2024). Findings of this study are consistent with the established understandings regarding the effect of drought stress on grain weight of rice.

### 3.4. Grain yield and dry matter partitioning

Upland rice varieties produced higher yields under non-stressed conditions (Figure 3a). However, grain yield significantly reduced under stressed conditions. Reduction in grain yields ranged 35-60% and 43-78% for short duration ranged 24-62 and 52-73% for long duration verities under moderately stressed and highly stressed conditions respectively (Figure 3a, Table S2). The results indicate that drought stress occurrence at later stages significantly influence upland rice yields. Lafitte et al. (Lafitte et al., 2004) reported 17-50% reduction in upland rice yield. Reduction in yield occurs as rice plant growth and development is affected under reduced water supply which is further linked with performance of yield components such as panicle number, number of spikelets, filled grain and grain weight. We observed in this study that performance of yield components such as tillering, panicle bearing capacity and grain weight was significantly affected under drought stress that potentially led to reduction in final reduction yields. This reduction in the rice yield under drought stress is consistent with the findings reported in research on drought stress evaluation in rice (Nokkoul and Wichitparp, 2014; Alou et al., 2018; Anyaoha et al., 2018; Hussain et al., 2018; da Mata et al., 2023; Pope et al., 2023; Sagar et al., 2025).

**Figure 3:**
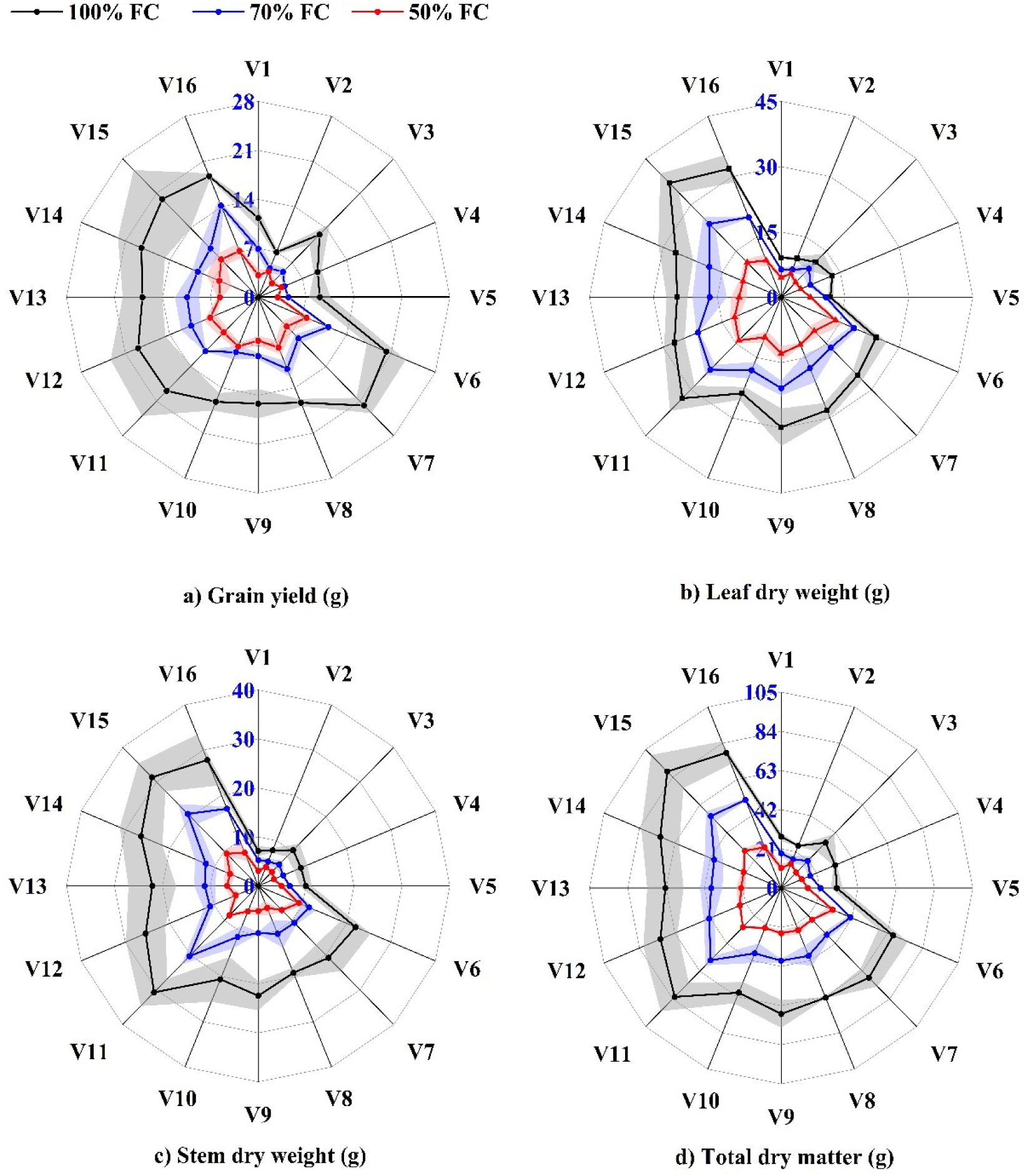
grain yield (GY), leaf dry weight (LDW), stem dry weight (SDW) and total dry matter (TDM) of sixteen upland varieties under non-stressed (100%), moderately stressed (70%) and highly stressed (50%) conditions (field capacity levels). Shaded areas show ± standard errors for means (*n*=3).

Dry matter partitioning of leaf (Figure 3b) and stem (Figure 3c) dry weights indicated that higher dry matter produced under non-stressed conditions. Leaf dry weights were reduced by 9-43% and 39-62% for short duration and by 22-38% and 43-71% for long duration upland rice varieties under moderately stressed and highly stressed conditions, respectively (Table S2). Stem dry weights also exhibited a similar trend with leaf dry weights, and the highest weights were observed under non-stressed conditions. Stem dry weights were reduced by 26-42% and 47-63% for short duration and by 34-57% and 58-80% for long duration upland rice varieties under moderately stressed and highly stressed conditions, respectively (Table S2). Total dry matter (Figure 3d) was higher under non-stress conditions, and it was significantly reduced under drought conditions. Reduction in total dry matter ranged 30-46% and 43-66% for short duration and by 34-48% and 54-70% for long duration upland rice varieties under moderately stressed and highly stressed conditions, respectively (Table S2). Growth and development of upland rice varieties is affected under drought stress as upland rice fails to extract water from soil layers. Although rice has greater root length, the ability of rice to extract water under drought stress is severely affected possibly due to factors like high soil water tension and the premature senescence of fine, absorptive root hairs, which collectively restrict water uptake from deeper soil profiles. Leaf, stem, and total dry matter of upland rice varieties are affected under water stress as key physiological processes are severely impacted (Dien et al., 2017; Jarin et al., 2024; Sutthachai et al., 2025). This reduction in dry matter accumulation is also linked with the plant’s growth stage (Melandri et al., 2021). Our results, showing significant dry matter loss are in line with the observations and findings of various published studies (Heinemann et al., 2015; Dien et al., 2017; Melandri et al., 2021; Yuwono et al., 2025).

### 3.5. Association among rice traits under non-stressed and drought-stressed conditions

Association among upland rice traits under non-stressed, moderately stressed and highly stressed conditions resulted in significant variations which indicate the drought stress influenced the performance of agronomic traits that led to variations in varietal performance (Figure 4). Phenological traits including days to flowering, days to first seed and days to maturity were highly associated (*p* < 0.001) under non-stressed conditions as well as drought stressed conditions. Drought stress delayed phenological development in most varieties. Significant positive association between phonological traits confirms that variety that flowered late, had more time to mature as well. Similar trend was observed by Ahmad et al. (Ahmad et al., 2020) in a field screening of rice germplasm. Plant height indicated highly significant (*p* < 0.001) association with days to flowering, days to first seed and days to maturity under non-stressed and moderately stressed conditions whereas no correlation was observed among these traits under highly stressed conditions. This indicated that plant heights possibly increase with longer growth duration under non stressed or moderately stressed conditions. Among the yield and yield traits, plant height was significantly positive correlated (*p* ≤ 0.05) with grain yield, grain weight, leaf and stem dry weights and total dry matter under non-stressed and moderately stressed conditions. This indicated that taller plants tended to accumulate higher dry matter and produced higher yield attributes. Catolos et al. (Catolos et al., 2017) found that plant height was clustered with biomass and growth rate of rice. However, we observed that under non-stressed conditions, plant height did not indicate significant correlation with number of tillers and panicles, though it was significantly (*p* ≤ 0.05) correlated with grain yield, grain weight, leaf and stem dry weight and total dry matter. These trends suggest that plant height possibly influenced yields primarily through dry matter accumulation and grain filling instead of tillering or panicle numbers. Number of tillers and panicles were highly significant (*p* < 0.001) and positively correlated which indicates higher more tillers produced more panicles. Generally, number of tillers or productive tillers and panicles are correlated and this algins with previous research (Lyu et al., 2014; Ahmad et al., 2020). Overall assessment indicated a strong association (*p* < 0.001) of number of tillers and panicles with grain yield, grain weight, leaf and stem dry weights and total dry matter however no significance was observed under applied treatments. This possibly resulted due to the experimental conditions where number of tillers and panicles generally did not influence the yield and yield components of assessed varieties. Grain yield indicated overall strong positive association (*p* < 0.001) with leaf and stem dry weights and total dry matter which indicated healthy and stronger plant produced higher yields. However, grain yield did not exhibit association with grain weight under non-stressed and moderately stressed conditions and exhibited a negative association under highly stressed conditions. Despite this grain yield was highly significant (*p* < 0.001) and positively correlated with leaf and stem dry weights and total dry matter under non-stressed, moderately stressed and highly stressed conditions. Overall grain weight indicated a positive association (*p* < 0.001) with leaf and stem dry weights and total dry matter, however it indicated a negative association (*p* < 0.05) with leaf dry weight and total dry matter under applied treatments which indicated that higher plant dry matter reduced grain weight possibly due to increased vegetative growth. Leaf and stem dry weights were highly significant (*p* < 0.001) and positively correlated under non-stressed, moderately stressed and highly stressed conditions and overall exhibited a significant positive association (*p* < 0.001). Supporting the results of this research, significant variable association among rice yield and yield contributing traits have been well documented in previous studies (Catolos et al., 2017; Ahmad et al., 2020; Khanal et al., 2022; Sarwendah et al., 2022; M et al., 2025; Sakran et al., 2025; Yuwono et al., 2025).

**Figure 4:**
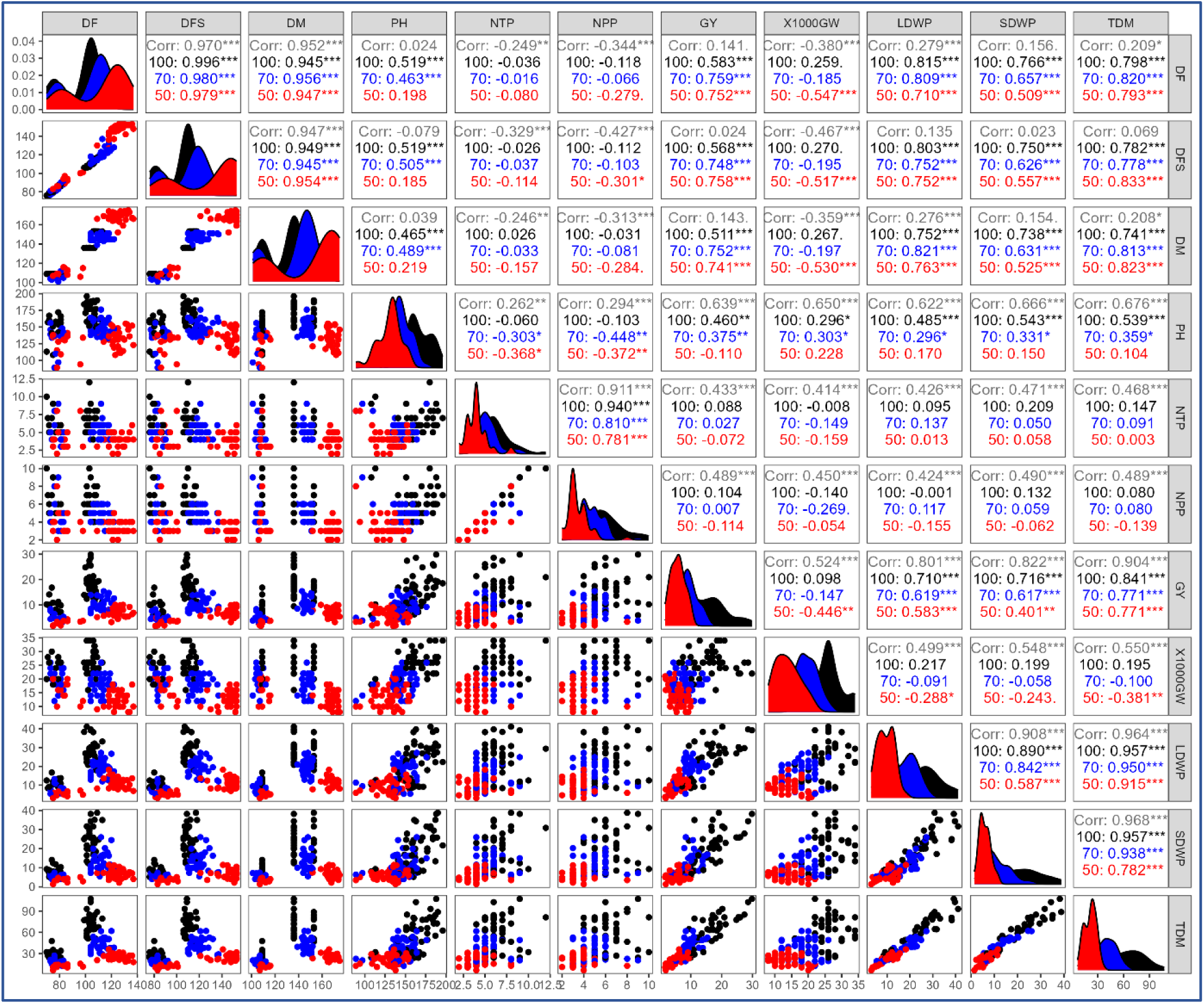
Correlation matrix, data distribution and scatter plots for phenology, growth, and yield traits of sixteen upland rice varieties under non stressed (100%), moderately stressed (70%) and highly stressed (50%) conditions (field capacity levels). Diagonals of the matrix indicate the distribution of each trait. Lower left of the diagonal indicates the scatter plots. Correlation values with significance levels presented with stars are shown on the upper right of the diagonal. DF: days for flowering, DFS: days to first seed, DM: days for maturity, PH: plant height, NTP: number of tillers per plant, NPP: number of panicles per plant, GY: grain yield, X1000: 1000 grain weight, LDWP: leaf dry weight per plant, SDWP: stem dry weight per plant, TDM: total dry matter, Highly significant at *p* < 0.001 (***), Moderately significant at *p* < 0.01 (**) and Significant at *p* < 0.05 (*).

### 3.6. Drought stress indices and varietal classification

Relative yield performance and drought stress indices were used to identify high yielding and drought stress resilient varieties (Table 2, 3). Under moderately stressed conditions, V6, V8, V11, V12, V14 and V16, exhibited higher relative yields compared to non-stress conditions and superior values for drought stress indices (Table 2). These varieties also indicated a linkage between them when evaluated using clustering (Figure 5). Though not superior in terms of yield performance but keeping in view the importance of early maturity and drought avoidance strategy, V1 and V3 exhibited higher relative yield and superior values for stress indices among short duration varieties (Table 2). In the hierarchical clustering these varieties exhibited linkage among other short duration varieties (Figure 5). Under highly stressed conditions, V6, V8, V10, V12 and V15 exhibited higher relative yields compared to non-stress conditions and superior values for drought stress indices (Table 3). Though not superior in terms of yield performance but keeping in view the importance of early maturity and drought avoidance strategy, V2 and V4 exhibited higher relative yield and superior values for stress indices among short duration varieties (Table 2). In the hierarchical clustering these varieties also exhibited linkage among other short duration varieties (Figure 5).

**Figure 5:**
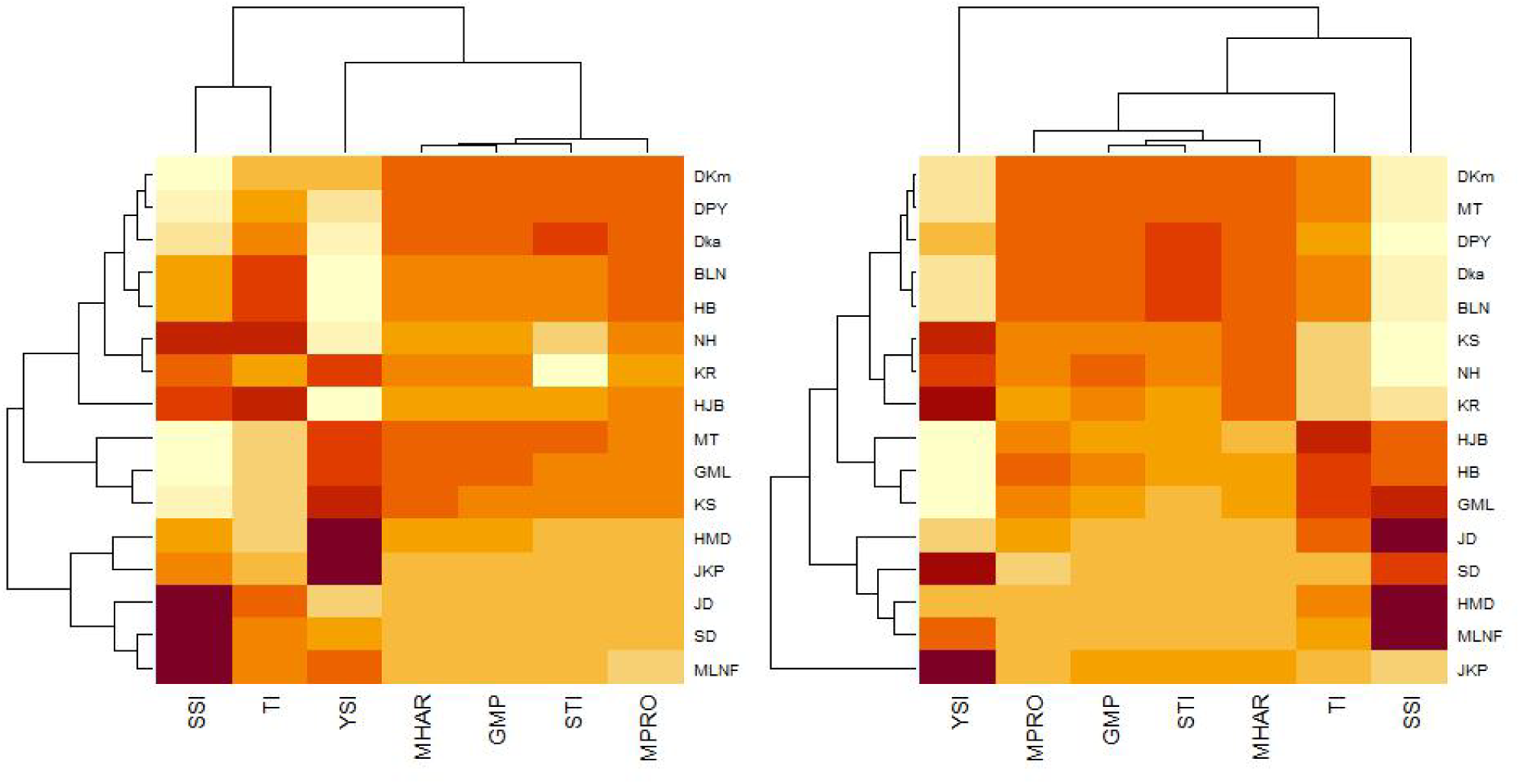
Heatmaps of drought stress indices among sixteen upland rice varieties under moderately stressed conditions at 70% field capacity (left) and highly stressed conditions at 50% field capacity (right). Intensity of the color in the figures indicates the degree of correlation among upland rice varieties and drought stress indices.

**Table 2.** Stress tolerance indices computed for sixteen upland rice varieties based on grain yield observed under non-stressed and moderately stressed conditions at 100% and 70% field capacity, respectively.

| Upland rice varieties | Yns | Yds | RYds | YSI | STI | M<sub>PRO</sub> | TI | GMP | SSI | M<sub>HAR</sub> |
| --- | --- | --- | --- | --- | --- | --- | --- | --- | --- | --- |
| 1 Hawm Mali Doi | 11.3 | 6.9 | 0.49 | 0.61 | 8.08 | 9.13 | 4.40 | 8.86 | 0.86 | 8.60 |
| 2 Jao Khao Pichit | 7.0 | 4.5 | 0.32 | 0.65 | 3.23 | 5.73 | 2.47 | 5.60 | 0.79 | 5.47 |
| 3 Jao Daeng | 12.7 | 5.1 | 0.36 | 0.40 | 6.65 | 8.88 | 7.57 | 8.04 | 1.33 | 7.27 |
| 4 Sahm Deuan | 9.4 | 4.2 | 0.29 | 0.44 | 4.03 | 6.78 | 5.23 | 6.26 | 1.24 | 5.77 |
| 5 Ma-led-nai-fai | 9.0 | 4.4 | 0.31 | 0.49 | 4.07 | 6.70 | 4.60 | 6.29 | 1.13 | 5.91 |
| 6 Dawk Kha | 20.3 | 11.1 | 0.78 | 0.55 | 23.07 | 15.67 | 9.20 | 14.98 | 1.01 | 14.32 |
| 7 Hawm Jet Ban | 21.9 | 8.3 | 0.59 | 0.38 | 18.67 | 15.08 | 13.57 | 13.47 | 1.38 | 12.03 |
| 8 Khao/ Sai | 16.3 | 11.1 | 0.78 | 0.68 | 18.56 | 13.68 | 5.23 | 13.43 | 0.71 | 13.18 |
| 9 Khao/ Ruang | 15.2 | 8.4 | 0.59 | 0.55 | 13.14 | 11.80 | 6.80 | 11.30 | 0.99 | 10.82 |
| 10 Nual Hawm | 16.2 | 8.5 | 0.60 | 0.53 | 14.14 | 12.33 | 7.67 | 11.72 | 1.05 | 11.14 |
| 11 Dawk Kham | 18.9 | 10.9 | 0.77 | 0.58 | 21.19 | 14.90 | 8.00 | 14.35 | 0.94 | 13.83 |
| 12 Dawk Pa-yawm | 19.0 | 10.6 | 0.75 | 0.56 | 20.72 | 14.80 | 8.40 | 14.19 | 0.98 | 13.61 |
| 13 Goo Meung Lung | 16.9 | 10.4 | 0.73 | 0.62 | 18.08 | 13.65 | 6.50 | 13.26 | 0.85 | 12.88 |
| 14 Hua Bon | 18.5 | 9.6 | 0.68 | 0.52 | 18.18 | 14.02 | 8.90 | 13.29 | 1.07 | 12.60 |
| 15 Bow Leb Nahag | 19.8 | 9.9 | 0.70 | 0.50 | 20.14 | 14.85 | 9.97 | 13.99 | 1.12 | 13.18 |
| 16 Mai Tahk | 18.7 | 14.2 | 1.00 | 0.76 | 27.30 | 16.45 | 4.57 | 16.29 | 0.54 | 16.13 |

**Table 3.**
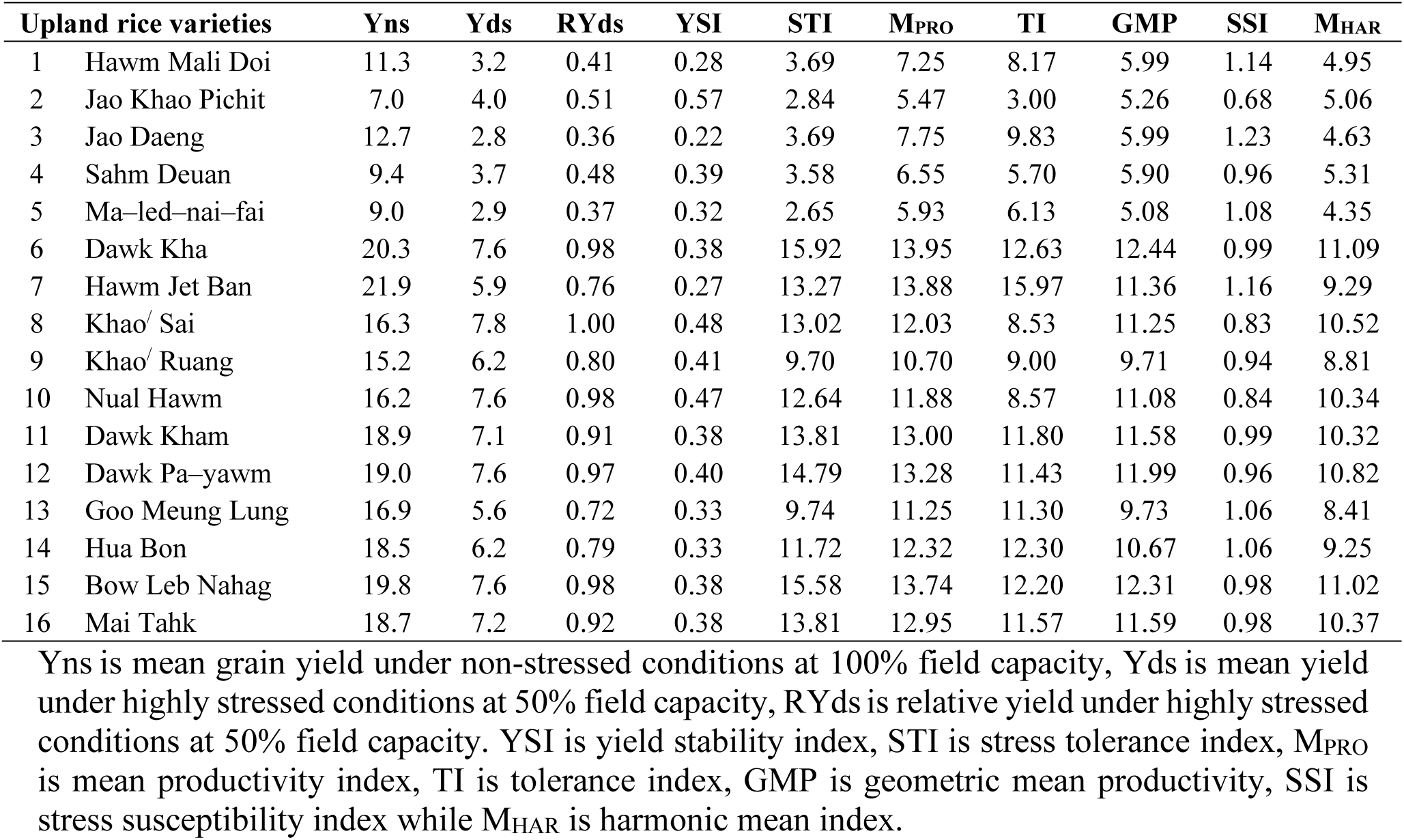
Stress tolerance indices computed for sixteen upland rice varieties based on grain yield observed under non-stressed and highly stressed conditions at 100% and 50% field capacity, respectively.

Yns is mean grain yield under non-stressed conditions at 100% field capacity, Yds is mean yield under moderately stressed conditions at 70% field capacity, RYds is relative yield under moderately stressed conditions at 70% field capacity. YSI is yield stability index, STI is stress tolerance index, M_PRO_ is mean productivity index, TI is tolerance index, GMP is geometric mean productivity, SSI is stress susceptibility index while M_HAR_ is harmonic mean index.

Yns is mean grain yield under non-stressed conditions at 100% field capacity, Yds is mean yield under highly stressed conditions at 50% field capacity, RYds is relative yield under highly stressed conditions at 50% field capacity. YSI is yield stability index, STI is stress tolerance index, M_PRO_ is mean productivity index, TI is tolerance index, GMP is geometric mean productivity, SSI is stress susceptibility index while M_HAR_ is harmonic mean index.

### 3.7. Association of Grain Yield with Drought Stress Indices

A strong positive (*p* < 0.001) association was observed between GY under non-stressed conditions, GY under moderately stressed conditions, GMP, STI, M_PRO_ and M_HAR_ (Figure 6). GMP, STI, M_PRO_, M_HAR_ and TI were strongly (*p* < 0.001) correlated with GY under non-stressed conditions. GMP, STI, M_PRO_ and M_HAR_ were strongly correlated (*p* < 0.001) with GY under moderately stressed conditions. YSI was significantly (*p* < 0.05) correlated whereas SSI indicated a significant (*p* < 0.05) negative association with GY under moderately stressed conditions (Figure 6). Both SSI and YSI were not correlated with GY under non-stressed conditions. There was a highly significant (*p* < 0.001) positive correlation among STI, GMP, M_PRO_ and M_HAR_ whereas a highly significant (*p* < 0.001) negative correlation was observed between SSI and YSI and a significant (*p* < 0.05) correlation between TI and YSI. However, TI and SSI were positively correlated (*p* < 0.05) (Figure 6).

**Figure 6:**
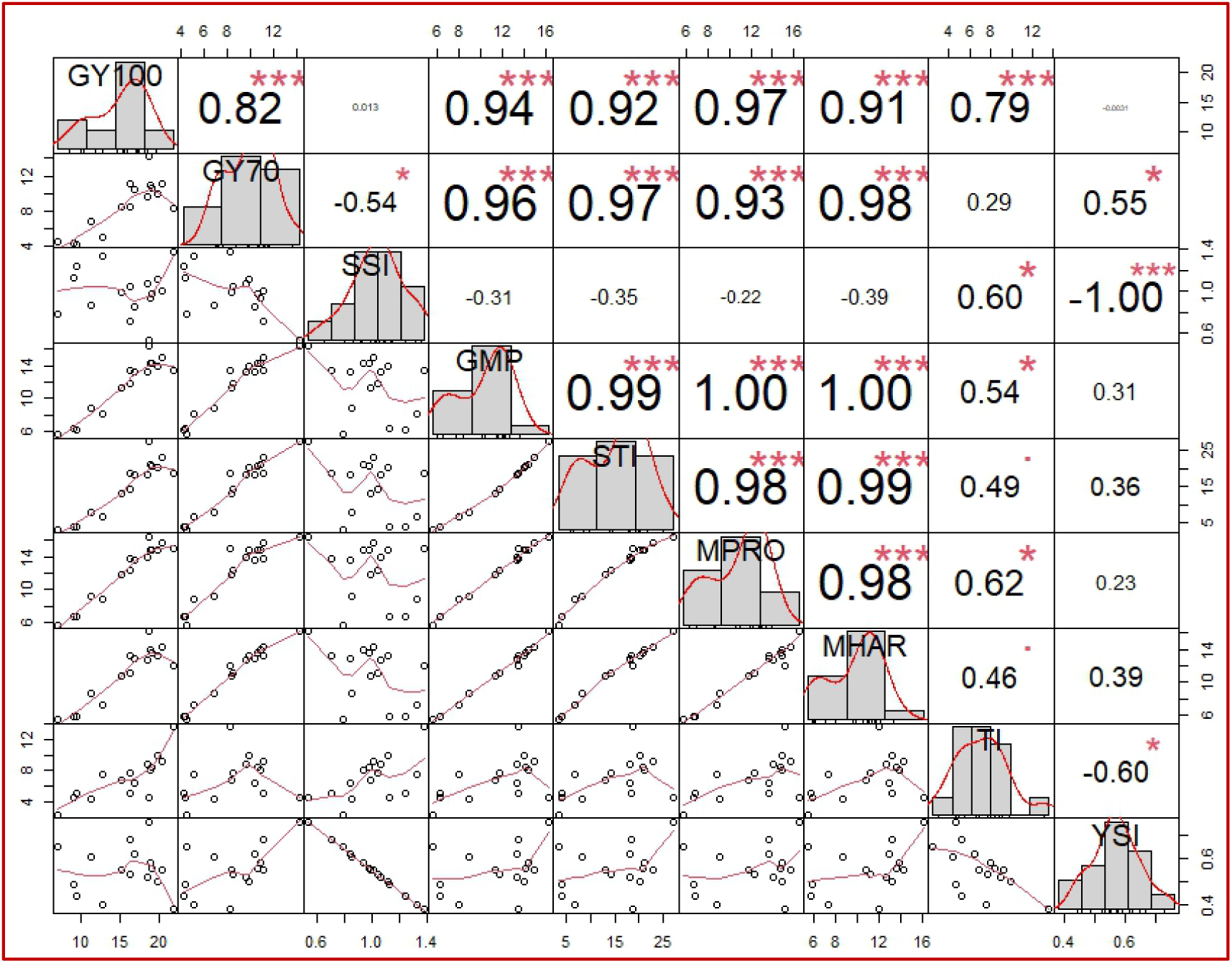
Pearson’s correlation matrix of grain yield under non-stressed conditions (GY100) at 100% field capacity (FC), grain yield under **moderately stressed conditions** (GY70) at 70% FC, stress susceptibility index (SSI), geometric mean productivity (GMP), stress tolerance index (STI), mean productivity index (M_PRO_), harmonic mean index (M_HAR_), tolerance index (TI) and yield stability index (YSI) for studied upland rice varieties. Diagonals of the matrix indicate the distribution of each. Lower left of the diagonal indicates the scatter plots with lines. Upper right of the diagonal indicates the correlation values and the corresponding significance. Intensity of color and the size of numerical values indicate the strength of correlation. Highly significant at *p* < 0.001 (***), Moderately significant at *p* < 0.01 (**) and Significant at *p* < 0.05 (*).

Under highly stressed conditions (Figure 7), strong positive (*p* < 0.001) association was observed between GY under non-stressed conditions, GY under highly stressed conditions, GMP, STI, M_PRO_ and M_HAR_. GMP, STI, M_PRO_, M_HAR_ and TI were strongly (*p* < 0.001) correlated with GY under non-stressed conditions. GMP, STI, M_PRO_ and M_HAR_ were strongly correlated (*p* < 0.001) with GY under highly stressed conditions. YSI and SSI were only two indices that did not indicate significant association with GY under non-stressed or highly stressed conditions (Figure 7). There was a highly significant (*p* < 0.001) positive correlation among STI, GMP, M_PRO_ and M_HAR_. TI was highly (*p* < 0.001) correlated with GMP, STI and M_PRO_, moderately (*p* < 0.001) correlated with M_HAR_ and correlated with SSI (*p* < 0.05). A highly significant (*p* < 0.001) and significant (*p* < 0.05) negative correlation was observed between SSI and YSI and TI and YSI, respectively (Figure 7). The effectiveness of the GMP, STI, M_PRO_ and M_HAR_ indices for screening rice genotypes under stress is well-documented (Raman et al., 2012; Wasae, 2021; Hallajian et al., 2024). Our findings align with previous research evidence, significantly demonstrating a positive correlation between these indices and grain yield under both moderate and high-stress environments. Debbarma et al. (Debbarma et al., 2024) identified tolerant cultivars using the TI, though the specific cultivars identified varied based on the individual trait analyzed. We observed that TI was not as effective and promising indices for grain yield as GMP, STI, M_PRO_ and M_HAR_ under both moderately and highly stressed conditions. Strong significant association (*p* < 0.001) of GMP, STI, M_PRO_, M_HAR_ with GY under non-stressed, moderately stressed and highly stressed conditions indicated that these indices were appropriate for their use as selection criteria for drought resilience when the conditions are moderately stressed. Under highly stressed conditions, TI also exhibited significant (*p* < 0.05) correlation with GY, there this could also be used.

**Figure 7:**
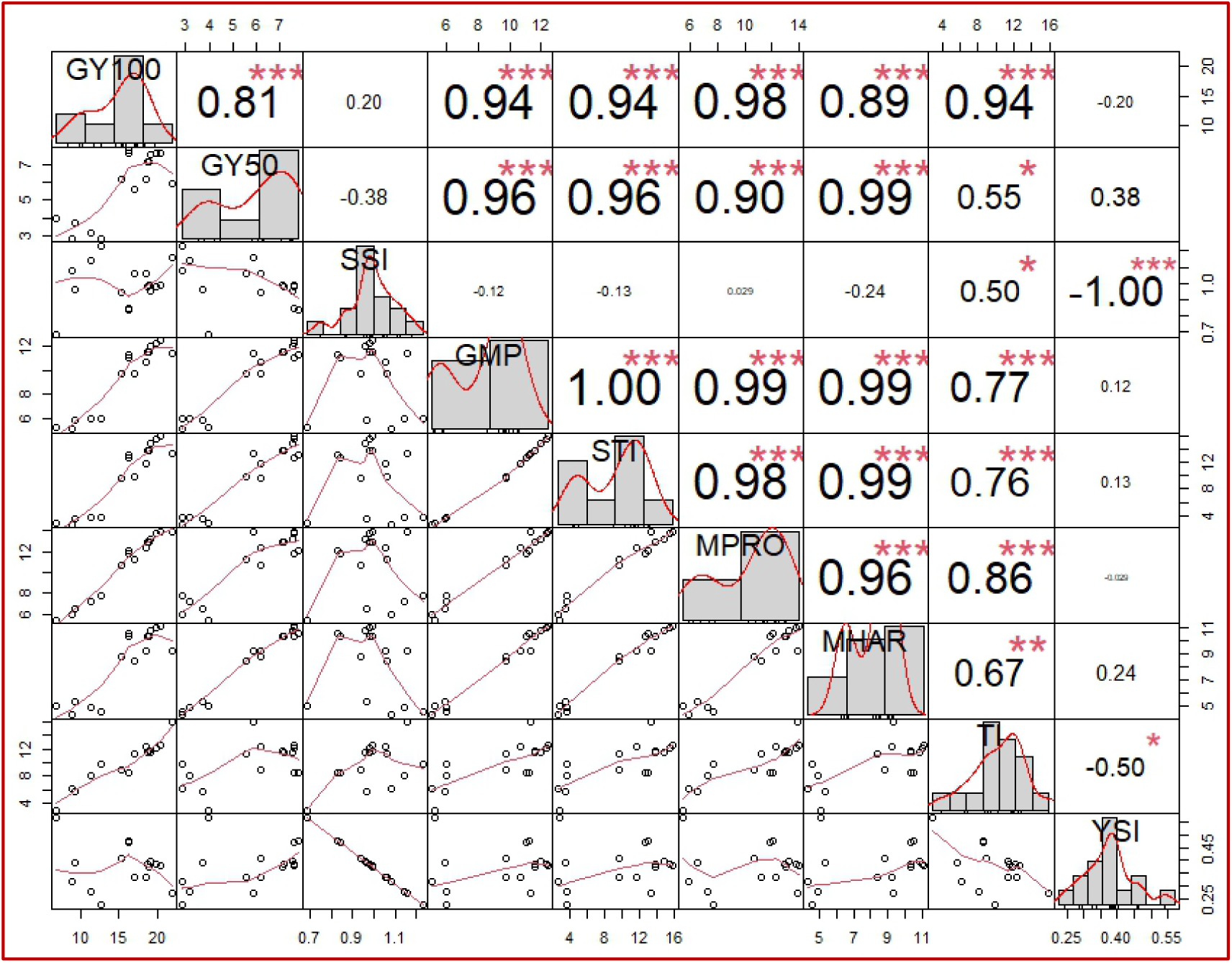
Pearson’s correlation matrix of grain yield under non-stressed conditions (GY100) at 100% field capacity (FC), grain yield under **highly stressed conditions** (GY50) at 50% FC, stress susceptibility index (SSI), geometric mean productivity (GMP), stress tolerance index (STI), mean productivity index (M_PRO_), harmonic mean index (M_HAR_), tolerance index (TI) and yield stability index (YSI) for studied upland rice varieties. Diagonals of the matrix indicate the distribution of each. Lower left of the diagonal indicates the scatter plots with lines. Upper right of the diagonal indicates the correlation values and the corresponding significance. Intensity of color and the size of numerical values indicate the strength of correlation. Highly significant at *p* < 0.001 (***), Moderately significant at *p* < 0.01 (**) and Significant at *p* < 0.05 (*).

## 4. Conclusion

Upland rice varieties were found unique in their agronomic responses and yield potential under non-stressed and stressed conditions. Short duration varieties particularly Ma-led-nai-fai (V5) indicated higher susceptibility to drought stress occurring at lateral crop stage. Short duration varieties other than V5 could be used for early planting and harvest. However, if environmental conditions are predicted to be not favourable at later stages growing long duration varieties may be advantageous. Among the long duration varieties, Dawk Kha (V6), Khao^/^ Sai (V8) and Dawk Pa–yawm (V12), exhibited superior performance, higher relative yields and promising values for stress indices under both moderately stressed and highly stressed conditions. Therefore, these varieties could be considered highly suitable for drought resilience and used to combat drought stress and maintain upland rice productivity. Under moderately stressed conditions varieties, Dawk Kham (V11), Goo Meung Lung (V13) and Mai Tahk (V16) and under highly stressed conditions varieties, Nual Hawm (V10) and Bow Leb Nahag (V15) exhibited better performance. These varieties could be used to obtain the desired traits in rice breeding. Ma-led-nai-fai (V5) from short duration varieties and Goo Meung Lung (V13) and Bow Leb Nahag (V15) could be used for acquiring traits for higher tillering and panicle bearing capacity. Short heighted varieties such as Jao Daeng (V3), Sahm Deuan (V4) and Ma-led-nai-fai (V5) could be used in breeding for short heighted new varieties to overcome lodging concerns. Overall varieties Dawk Kha (V6), Khao^/^ Sai (V8) and Dawk Pa–yawm (V12), indicated higher stability under stressed conditions therefore, these long duration varieties could be used for obtaining better yields under diverse agroclimatic conditions and under unpredicted weather patterns whereas other promising varieties indicating diversity could be used for specific traits in breeding program. Strong significant association of GMP, STI, M_PRO_, M_HAR_ with grain yield under non-stressed, moderately stressed and highly stressed conditions indicated that these indices were appropriate for their use as selection criteria for drought resilience.

## Author Contributions

T.H: Conceptualization, funding acquisition, methodology, investigation, data collection, data curation, formal analysis, visualization, writing – original draft. J.A: Conceptualization, methodology, supervision, project management, resources, writing – review and editing. C.N: Supervision, resources, technical support, writing – review and editing. A.A: Methodology, data collection, writing – review and editing. T.K: writing – review and editing.

## Acknowledgement

The authors acknowledge Graduate School, Prince of Songkla University, Thailand and the Faculty of Natural Resources, Prince of Songkla University, Thailand for providing financial support and research resources. Research was supported by Higher Education Research Promotion and the Thailand’s Education Hub for Southern Region of ASEAN Countries, Project Office of the Higher Education Commission.

## References

Adeboye, K. A., Oduwaye, O. A., Daniel, I. O., Fofana, M., and Semon, M. (2021). Characterization of flowering time response among recombinant inbred lines of WAB638-1/PRIMAVERA rice under reproductive stage drought stress. Plant Genet. Resour. Charact. Util. 19, 1–8. doi: DOI: 10.1017/S1479262121000010.

Ahmad, M. S., Wu, B., Wang, H., and Kang, D. (2020). Field Screening of Rice Germplasm (Oryza sativa L. ssp. japonica) Based on Days to Flowering for Drought Escape. Plants 9. doi: 10.3390/plants9050609.

Alou, I. N., Steyn, J. M., Annandale, J. G., and van der Laan, M. (2018). Growth, phenological, and yield response of upland rice (Oryza sativa L. cv. Nerica 4®) to water stress during different growth stages. Agric. Water Manag. 198, 39–52. doi: 10.1016/j.agwat.2017.12.005.

Anwar, J., Subhani, G. M., Hussain, M., Ahmad, J., Hussain, M., and Munir, M. (2011). Drought tolerance indices and their correlation with yield in exotic wheat genotypes. Pakistan J. Bot. 43, 1527–1530.

Anyaoha, C. O., Fofana, M., Gracen, V. E., Tongoona, P. B., Blay, E. T., Semon, M., et al. (2018). Yield potential of upland rice varieties under reproductive-stage drought and optimal water regimes in Nigeria. Plant Genet. Resour. Charact. Util. 16, 378–385. doi: DOI: 10.1017/S1479262118000060.

Arif, A., Parveen, N., Waheed, M. Q., Atif, R. M., Waqar, I., and Shah, T. M. (2021). A Comparative Study for Assessing the Drought-Tolerance of Chickpea Under Varying Natural Growth Environments. Front. Plant Sci. 11. doi: 10.3389/fpls.2020.607869.

Bhandari, U., Gajurel, A., Khadka, B., Thapa, I., Chand, I., Bhatta, D., et al. (2023). Morpho-physiological and biochemical response of rice (Oryza sativa L.) to drought stress: A review. Heliyon 9, e13744. doi: 10.1016/j.heliyon.2023.e13744.

Bouslama, M., and Schapaugh, W. T. (1984). Stress tolerance in soybeans. I. Evaluation of three screening techniques for heat and drought tolerance. Crop Sci. 24, 933–937.

Catolos, M., Sandhu, N., Dixit, S., and Shamsudin, N. A. A. (2017). Genetic Loci Governing Grain Yield and Root Development under Variable Rice Cultivation Conditions. 8, 1–17. doi: 10.3389/fpls.2017.01763.

Chatterjee, A., Galiba, G., Kocsy, G., Kar, R. K., and Dey, N. (2024). Molecular insight into drought tolerance of CR Dhan 40: an upland rice line from Eastern India. J. Crop Sci. Biotechnol. 27, 225–234. doi: 10.1007/s12892-023-00222-3.

Chen, X., Xiao, D., Qi, Y., Shi, Z., Bai, H., Lu, Y., et al. (2025). Projected future changes in extreme climate indices affecting rice production in China using a multi-model ensemble of CMIP6 projections. Front. Plant Sci. 16, 1595367. doi: 10.3389/fpls.2025.1595367.

Clarke, J. M., Townley-Smith, F., McCaig, T. N., and Green, D. G. (1984). Growth analysis of spring wheat cultivars of varying drought resistance 1. Crop Sci. 24, 537–541.

da Mata, C. R., de Castro, A. P., Lanna, A. C., Bortolini, J. C., and de Moraes, M. G. (2023). Physiological and yield responses of contrasting upland rice genotypes towards induced drought. Physiol. Mol. Biol. Plants 29, 305–317. doi: 10.1007/s12298-023-01287-8.

Davatgar, N., Neishabouri, M. R., Sepaskhah, A. R., and Soltani, A. (2012). Physiological and morphological responses of rice (Oryza sativa L.) to varying water stress management strategies. Int. J. Plant Prod. 3, 19–32. doi: 10.22069/ijpp.2012.660.

Debbarma, S., Roy, M., Nayak, J. K., Poddar, S., Saha, P., Sharma, P., et al. (2024). Rapid Screening of Traditional Rice Cultivars against Drought Using Tolerance Indices of Seedlings. Int. J. Environ. Clim. Chang. 14, 352–369.

Dien, D. C., Yamakawa, T., Mochizuki, T., and Htwe, A. Z. (2017). Dry Weight Accumulation, Root Plasticity, and Stomatal Conductance in Rice (Oryza sativa L.) Varieties under Drought Stress and Re-Watering Conditions. 3189–3206. doi: 10.4236/ajps.2017.812215.

El-Refaee, Y. Z., Seadh, S. E., Abdel-Moneam, M. A., and Eltantawy, M. E. M. (2023). Determination of Drought Tolerance Indices as Selection Criteria of Rice Genotypes under Water Deficit Conditions in Egypt. Int. J. Plant Soil Sci. 35, 192–208.

Fernandez, G. C. (1992). Effective selection criteria for assessing plant stress tolerance. in Proceeding of the International Symposium on Adaptation of Vegetables and other Food Crops in Temperature and Water Stress (13–16 August 1992, Shanhua, Taiwan), 257–270.

Fischer, R. A., and Maurer, R. (1978). Drought resistance in spring wheat cultivars. I. Grain yield responses. Aust. J. Agric. Res. 29, 897–912.

Fu, J., Jian, Y., Wang, X., Li, L., Ciais, P., Zscheischler, J., et al. (2023). Extreme rainfall reduces one-twelfth of China’s rice yield over the last two decades. Nat. food 4, 416–426. doi: 10.1038/s43016-023-00753-6.

Golabadi, M., Arzani, A., and Maibody, M. (2006). Assessment of Drought Tolerance in Segregating Populations in Durum Wheat. African J. Agric. Res. 1, 162–171.

Habib, M. A., Azam, M. G., Haque, M. A., Hassan, L., Khatun, M. S., Nayak, S., et al. (2024). Climate-smart rice (Oryza sativa L.) genotypes identification using stability analysis, multi-trait selection index, and genotype-environment interaction at different irrigation regimes with adaptation to universal warming. Sci. Rep. 14, 13836. doi: 10.1038/s41598-024-64808-9.

Hallajian, M. T., Ebadi, A. A., and Kordrostami, M. (2024). Advancing rice breeding for drought tolerance: a comprehensive study of traditional and mutant lines through agronomic performance and drought tolerance indices. BMC Plant Biol. 24, 1087. doi: 10.1186/s12870-024-05771-5.

Heinemann, A. B., Barrios-Perez, C., Ramirez-Villegas, J., Arango-Londoño, D., Bonilla-Findji, O., Medeiros, J. C., et al. (2015). Variation and impact of drought-stress patterns across upland rice target population of environments in Brazil. J. Exp. Bot. 66, 3625–3638. doi: 10.1093/jxb/erv126.

Hossain, A. B. S., Sears, R. G., Cox, T. S., and Paulsen, G. M. (1990). Desiccation tolerance and its relationship to assimilate partitioning in winter wheat. Crop Sci. 30, 622–627.

Hussain, N., Ahmed, M., Duangpan, S., Hussain, T., and Taweekun, J. (2021a). Potential impacts of water stress on rice biomass composition and feedstock availability for bioenergy production. Sustainability 13, 10449.

Hussain, T. (2017). Modeling Upland Rice for Drought Tolerance Using DSSAT.

Hussain, T., Anothai, J., Nualsri, C., Ata-Ul-Karim, S. T., Duangpan, S., Hussain, N., et al. (2023). Assessment of CSM–CERES–Rice as a Decision Support Tool in the Identification of High-Yielding Drought-Tolerant Upland Rice Genotypes. Agronomy 13, 432.

Hussain, T., Anothai, J., Nualsri, C., and Soonsuwon, W. (2018). Application of CSM-CERES-Rice in scheduling irrigation and simulating effect of drought stress on upland rice yield. Indian J. Agric. Res. 52, 140–145.

Hussain, T., Hussain, N., Ahmed, M., Nualsri, C., and Duangpan, S. (2021b). Responses of lowland rice genotypes under terminal water stress and identification of drought tolerance to stabilize rice productivity in southern Thailand. Plants 10, 2565.

Ichsan, C. N., Basyah, B., Zakaria, S., and Efendi, E. (2020). Differences of water status and relationship with roots growth and yield of rice under water stress. Syst. Rev. Pharm. 11, 611–618.

IRRI (2014). Standard Evaluation System for Rice. 5th ed. Los Banos, the Philippines International Rice Research Institute.

Jarin, A. S., Islam, M. M., Rahat, A., Ahmed, S., Ghosh, P., and Murata, Y. (2024). Drought Stress Tolerance in Rice: Physiological and Biochemical Insights. Int. J. Plant Biol. 15, 692–718. doi: 10.3390/ijpb15030051.

Jiang, X., Li, G., and Luo, S. (2025). The research progress on the mechanisms of drought stress at different growth stages affecting rice yield formation. Resour. Data J. 4, 302–321. doi: 10.50908/rdj.4.0_302.

Joseph, M., Moonsammy, S., Davis, H., Warner, D., Adams, A., and Timothy Oyedotun, T. D. (2023). Modelling climate variabilities and global rice production: A panel regression and time series analysis. Heliyon 9. doi: 10.1016/j.heliyon.2023.e15480.

Kang, D.-J., and Futakuchi, K. (2019). Effect of Moderate Drought-Stress on Flowering Time of Interspecific Hybrid Progenies (Oryza sativa L. × Oryza glaberrima Steud.). J. Crop Sci. Biotechnol. 22, 75–81. doi: 10.1007/s12892-019-0015-0.

Khanal, P., Bigyan, K. C., Lamichhane, S., Raj, N., Sharma, S., and Upadhyay, K. (2022). CORRELATION AND PATH ANALYSIS OF YIELD AND YIELD ATTRIBUTING CHARACTERS IN RICE UNDER REPRODUCTIVE DROUGHT STRESS CONDITION. Rev. Food Agric. 3, 39–42. doi: 10.26480/rfna.01.2022.39.42.

Khotasena, S., Sanitchon, J., Chankaew, S., and Monkham, T. (2022). The Basic Vegetative Phase and Photoperiod Sensitivity Index as the Major Criteria for Indigenous Upland Rice Production in Thailand under Unpredictable Conditions. Agronomy 12. doi: 10.3390/agronomy12040957.

Konaté, A. K., Zongo, A., Sangaré, J. R., Dardou, A., and Audebert, A. (2022). Effect of water stress on growth, yield and yield components of rice (Oryza sativa L.) genotypes. Int. J. Sci. Res. Arch 5, 28–38. doi: 10.30574/ijsra.2022.5.1.0030.

Lafitte, H. R., Price, A. H., and Courtois, B. (2004). Yield response to water deficit in an upland rice mapping population: associations among traits and genetic markers. Theor. Appl. Genet. 109, 1237–1246. doi: 10.1007/s00122-004-1731-8.

Lanna, A., Costa Coelho, G., Souza Moreira, A., Rios Terra, T., Brondani, C., Rios Saraiva, G., et al. (2021). Upland rice: phenotypic diversity for drought tolerance. Sci. Agric. 78. doi: 10.1590/1678-992x-2019-0338.

Liberatore, C. M., Biancucci, M., Ezquer, I., Gregis, V., and Di Marzo, M. (2025). Investigating how reproductive traits in rice respond to abiotic stress. J. Exp. Bot. 76, 2064–2080. doi: 10.1093/jxb/eraf031.

Liu, Z., Lv, A., and Li, T. (2025). Intensified Drought Threatens Future Food Security in Major Food-Producing Countries. Atmosphere (Basel). 16. doi: 10.3390/atmos16010034.

Lyu, J., Li, B., He, W., Zhang, S., Gou, Z., Zhang, J., et al. (2014). A genomic perspective on the important genetic mechanisms of upland adaptation of rice. BMC Plant Biol. 14, 160. doi: 10.1186/1471-2229-14-160.

M, A., M, V. P., N, R. S. R., B, K. S., M, V. R., and G, S. B. (2025). Association Analysis and Correlation Studies in Upland Rice (Oryza sativa). Int. J. Plant Soil Sci. 37, 46–53. doi: 10.9734/ijpss/2025/v37i55428.

Mansour, E., Desoky, E. S. M., Ali, M. M. A., Abdul-Hamid, M. I., Ullah, H., Attia, A., et al. (2021). Identifying drought-tolerant genotypes of faba bean and their agro-physiological responses to different water regimes in an arid Mediterranean environment. Agric. Water Manag. 247, 106754. doi: 10.1016/j.agwat.2021.106754.

Melandri, G., AbdElgawad, H., Floková, K., Jamar, D. C., Asard, H., Beemster, G. T. S., et al. (2021). Drought tolerance in selected aerobic and upland rice varieties is driven by different metabolic and antioxidative responses. Planta 254, 13. doi: 10.1007/s00425-021-03659-4.

Nokkoul, R., and Wichitparp, T. (2014). EFFECT OF DROUGHT CONDITION ON GROWTH, YIELD AND GRAIN QUALITY OF UPLAND RICE. Am. J. Agric. Biol. Sci. 9. doi: 10.3844/ajabssp.2014.439.444.

Pope, E. M., Opile, W., Ngode, L., and Chepkoech, E. (2023). Effect of Water Stress Duration on the Growth Characteristics and Yield Components of Upland Rice Varieties in Kenya. Asian J. Res. Crop Sci. 8, 273–286. doi: 10.9734/ajrcs/2023/v8i4208.

R Core Team (2024). R: A Language and Environment for Statistical Computing. Available at: https://www.r-project.org/.

Rahman, M. T., Islam, M. T., and Islam, M. O. (2002). Effect of water stress at different growth stages on yield and yield contributing characters of transplanted Aman rice. Pakistan J. Biol. Sci. 5, 169–172. doi: 10.3923/pjbs.2002.169.172.

Raman, A., Verulkar, S. B., Mandal, N. P., Variar, M., Shukla, V. D., Dwivedi, J. L., et al. (2012). Drought yield index to select high yielding rice lines under different drought stress severities. Rice 5, 1–12. doi: 10.1186/1939-8433-5-31.

Rashid, A., Saleem, Q., Nazir, A., and Kazım, H. S. (2003). Yield potential and stability of nine wheat varieties under water stress conditions. Int. J. Agric. Biol. 5, 7–9.

Rehmani, M. I. A., Ding, C., Li, G., Ata-Ul-Karim, S. T., Hadifa, A., Bashir, M. A., et al. (2021). Vulnerability of rice production to temperature extremes during rice reproductive stage in Yangtze River Valley, China. J. King Saud Univ. - Sci. 33, 101599. doi: 10.1016/j.jksus.2021.101599.

Rosielle, A. A., and Hamblin, J. (1981). Theoretical aspects of selection for yield in stress and non-stress environments. Crop Sci. 21, 943–946.

Sadhukhan, D., Mukherjee, T., Sarkar, A., Devi, N. D., Bisarya, D., Kumar, V., et al. (2024). A Comprehensive Analysis of Drought Stress Responses in Rice (Oryza sativa L.): Insights into Developmental Stage Variations from Germination to Grain Filling. Int. J. Environ. Clim. Chang. 14, 141–158.

Sagar, S., Ramamoorthy, P., Ramalingam, S., Muthurajan, R., Natarajan, S., Doraiswamy, U., et al. (2025). Drought’s physiological footprint: implications for crop improvement in rice. Mol. Biol. Rep. 52, 298. doi: 10.1007/s11033-025-10405-6.

Sakran, R. M., Ghazy, M. I., Gaballah, M. M., Hussein, F. A., Aamer, S. M., Ghazy, H. A., et al. (2025). Genetic variability in physiological and agronomic traits of newly developed rice lines under well-watered and water-deficit conditions. BMC Plant Biol. 25, 1291. doi: 10.1186/s12870-025-07436-3.

Sarwendah, M., Lubis, I., Junaedi, A., and Purwoko, B. S. (2022). Application of selection index for rice mutant screening under a drought stress condition imposed at reproductive growth phase. BIODIVERSITAS 23, 5446–5452. doi: 10.13057/biodiv/d231056.

Schneider, K. A., Rosales-serna, R., Ibarra-perez, F., Cazares-enriquez, B., Acosta-gallegos, J. A., Ramirez-vallejo, P., et al. (1997). Improving common bean performance under drought stress. Crop Sci. 37, 43–50.

Singh, B., Reddy, K. R., Redoña, E. D., and Walker, T. (2017). Screening of Rice Cultivars for Morpho-Physiological Responses to Early-Season Soil Moisture Stress. Rice Sci. 24, 322–335. doi: 10.1016/j.rsci.2017.10.001.

Sun, X., Xiong, H., Jiang, C., Zhang, D., Yang, Z., Huang, Y., et al. (2022). Natural variation of DROT1 confers drought adaptation in upland rice. Nat. Commun. 13, 4265. doi: 10.1038/s41467-022-31844-w.

Sutthachai, S., Trunjaruen, A., Mahatthanaphatcharakun, P., and Taratima, W. (2025). Physio-biochemical and anatomical responses of upland rice (Oryza sativa L.) genotype during the vegetative stage under drought stress. Asian J. Agric. Biol. 2025, 2024265.

Todaka, D., Shinozaki, K., and Yamaguchi-shinozaki, K. (2015). Recent advances in the dissection of drought-stress regulatory networks and strategies for development of drought-tolerant transgenic rice plants. 6, 1–20. doi: 10.3389/fpls.2015.00084.

Wasae, A. (2021). Evaluation of Drought Stress Tolerance Based on Selection Indices in Haricot Bean Varieties Exposed to Stress at Different Growth Stages. Int. J. Agron. 2021. doi: 10.1155/2021/6617874.

Wopereis, M. C. S., Kropff, M. J., Maligaya, A. R., and Tuong, T. P. (1996). Drought-stress responses of two lowland rice cultivars to soil water status. F. Crop. Res. 46, 21–39. doi: 10.1016/0378-4290(95)00084-4.

Yang, X., Liu, C., Niu, X., Wang, L., Li, L., Yuan, Q., et al. (2022). Research on lncRNA related to drought resistance of Shanlan upland rice. BMC Genomics 23, 336. doi: 10.1186/s12864-022-08546-0.

Yuwono, S. S., Ghulamahdi, M., and Palupi, E. R. (2025). Morphological, physiological and rice yield in lowland and upland under drought stress. Aust. J. Crop Sci. 19, 398–407.

Zain, N. A. M., Ismail, M. R., Puteh, A., Mahmood, M., and Islam, M. R. (2014). Impact of cyclic water stress on growth, physiological responses and yield of rice (Oryza sativa L.) grown in tropical environment. Ciência Rural 44, 2136–2141. doi: 10.1590/0103-8478cr20131154.

Zhang, C., Liu, J., Zhao, T., Gomez, A., Li, C., Yu, C., et al. (2016). A Drought-Inducible Transcription Factor Delays Reproductive Timing in Rice. Plant Physiol. 171, 334–343. doi: 10.1104/pp.16.01691.

Zulkarnain, W. M., Ismail, M. R., Saud, H. M., Othman, R., Habib, S. H., and Kausar, H. (2013). Growth and yield response to water availability at different growth stages of rice. J. Food, Agric. Environ. 11, 540–544.

