## Supplementary Table 1 and 2 for "Assessing Drought Resilience and Identification of High Yielding Upland Rice Varieties through Phenology, Growth and Yield Traits"

### Supplementary Tables

**Table S1:** Changes in performance of days to flowering, days to first seed, days to maturity, plant height, number of tillers and number of panicles of sixteen upland rice varieties under non-stressed (100%), moderately stressed (70%) and highly stressed (50%) conditions (field capacity levels).

| Upland Rice Varieties | DF | | DS | | DM | | PH | | NT | | NP | |
|---|---|---|---|---|---|---|---|---|---|---|---|---|
| | days | | days | | days | | % | | no. | | no. | |
| | 70 | 50 | 70 | 50 | 70 | 50 | 70 | 50 | 70 | 50 | 70 | 50 |
| Hawm Mali Doi | 1 | 4 | 2 | 6 | -5 | -2 | -12 | -20 | -1 | -2 | -1 | -2 |
| Jao Khao Pichit | 2 | 5 | 2 | 5 | -2 | 1 | -9 | -15 | -1 | -1 | -1 | -2 |
| Jao Daeng | 2 | 5 | 2 | 6 | -3 | 1 | -9 | -15 | -2 | -3 | -2 | -3 |
| Sahm Deuan | 2 | 9 | 2 | 13 | -1 | 6 | -10 | -13 | -1 | -2 | -2 | -2 |
| Ma–led–nai–fai | 4 | 12 | 4 | 13 | -5 | -1 | -20 | -22 | -1 | -2 | -2 | -4 |
| Dawk Kha | 7 | 17 | 7 | 35 | -3 | 15 | -8 | -22 | -1 | -2 | -1 | -3 |
| Hawm Jet Ban | 9 | 25 | 9 | 40 | 11 | 27 | -9 | -18 | 0 | -1 | 0 | -2 |
| Khao/ Sai | 9 | 19 | 11 | 40 | 14 | 35 | -12 | -22 | -1 | -2 | -1 | -2 |
| Khao/ Ruang | 7 | 21 | 8 | 42 | 13 | 33 | -21 | -30 | -1 | -3 | -1 | -3 |
| Nual Hawm | 3 | 19 | 4 | 32 | 9 | 32 | -17 | -25 | -2 | -3 | -2 | -3 |
| Dawk Kham | 5 | 14 | 5 | 34 | -3 | 18 | -9 | -15 | 0 | -2 | 0 | -3 |
| Dawk Pa–yawm | 5 | 15 | 5 | 24 | 5 | 29 | -12 | -21 | -1 | -2 | -1 | -2 |
| Goo Meung Lung | 8 | 22 | 14 | 31 | 6 | 24 | -10 | -18 | -2 | -5 | -4 | -5 |
| Hua Bon | 7 | 20 | 14 | 37 | 0 | 26 | -15 | -25 | -1 | -2 | -1 | -3 |
| Bow Leb Nahag | 9 | 24 | 14 | 39 | 14 | 36 | -14 | -28 | -3 | -4 | -3 | -4 |
| Mai Tahk | 10 | 25 | 13 | 38 | -1 | 21 | -13 | -26 | -2 | -3 | -1 | -3 |

Changes in days to flowering (DF), days to first seed (DS) and days to maturity (DM) are presented by difference in number of days. Number of tillers (NT) and number of panicles (NP) are presented by difference in numbers (no.), whereas changes in plant height (PH), are presented by % difference.

**Table S2:** Changes in performance of grain weight, grain yield, leaf dry weight, stem dry weight and total dry matter under non-stressed (100%), moderately stressed (70%) and highly stressed (50%) conditions (field capacity levels).

| **Upland Rice Varieties** | **1000GW** | | **GY** | | **LDW** | | **SDW** | | **TDM** | |
|---|---|---|---|---|---|---|---|---|---|---|
| | **%** | | | | | | | | | |
| | **70** | **50** | **70** | **50** | **70** | **50** | **70** | **50** | **70** | **50** |
| Hawm Mali Doi | -15 | -23 | -39 | -72 | -30 | -50 | -26 | -57 | -32 | -61 |
| Jao Khao Pichit | -13 | -28 | -35 | -43 | -29 | -39 | -32 | -47 | -32 | -43 |
| Jao Daeng | -19 | -35 | -60 | -78 | -19 | -58 | -40 | -61 | -40 | -66 |
| Sahm Deuan | -19 | -35 | -56 | -61 | -43 | -62 | -42 | -63 | -46 | -62 |
| Ma–led–nai–fai | -21 | -36 | -51 | -68 | -9 | -40 | -34 | -51 | -30 | -52 |
| Dawk Kha | -32 | -55 | -45 | -62 | -23 | -43 | -48 | -58 | -38 | -54 |
| Hawm Jet Ban | -22 | -46 | -62 | -73 | -35 | -57 | -49 | -66 | -48 | -65 |
| Khao/ Sai | -29 | -48 | -32 | -52 | -37 | -58 | -45 | -75 | -38 | -62 |
| Khao/ Ruang | -26 | -50 | -45 | -59 | -30 | -57 | -57 | -77 | -42 | -64 |
| Nual Hawm | -30 | -53 | -47 | -53 | -24 | -58 | -46 | -73 | -38 | -62 |
| Dawk Kham | -31 | -50 | -42 | -62 | -28 | -57 | -34 | -72 | -34 | -64 |
| Dawk Pa–yawm | -34 | -52 | -44 | -60 | -22 | -57 | -57 | -80 | -40 | -66 |
| Goo Meung Lung | -39 | -58 | -38 | -67 | -31 | -60 | -49 | -71 | -40 | -65 |
| Hua Bon | -40 | -57 | -48 | -67 | -32 | -64 | -55 | -76 | -45 | -69 |
| Bow Leb Nahag | -33 | -53 | -50 | -62 | -36 | -70 | -34 | -70 | -38 | -68 |
| Mai Tahk | -31 | -57 | -24 | -62 | -38 | -71 | -39 | -74 | -35 | -70 |

Changes in 1000 grain weight (1000GW), grain yield (GY), leaf dry weight (LDW), stem dry weight (SDW) and total dry matter (TDM) are presented by % difference.
